# Early-life core microbiota are associated with postlarval survival of *Penaeus vannamei*: A meta-analysis

**DOI:** 10.64898/2026.10.22.760395

**Authors:** Rudan Chen, Weiyue Zhang, Junqi Yu, Haoyu Song, Yanshuo Zhou, Chen Chen, Xianke Huang, Demin Zhang, Maocang Yan, Kai Wang

## Abstract

Larviculture is critical to the production of *Penaeus vannamei*, the world’s leading farmed shrimp species, but remains constrained by erratic postlarval survival. Although early-life microbiome establishment is increasingly recognized as essential for larval health and survival, microbiome-based hatchery interventions are hindered by limited understanding of larval core microbiota. Here, we synthesized publicly available and self-generated 16S rRNA gene amplicon datasets of larval and rearing-water microbiota from 13 studies encompassing seven major producing regions worldwide to characterize shrimp larval core microbiota, their assembly processes, and their associations with larval survival. Although larval prokaryotic communities exhibited stronger geographic variation than developmental-stage succession, a conserved set of dominant bacterial families, including *Roseobacteraceae*, *Vibrionaceae*, *Flavobacteriaceae*, *Moraxellaceae*, and *Pseudoalteromonadaceae*, consistently prevailed across developmental stages and regions. By integrating occupancy, persistence, and ecological network centrality analyses, we demonstrated that common, temporal, and ecological core microbiota differ substantially in composition, diversity, stage specificity, and ecological roles in network stability. Compared with non-core taxa, core taxa were generally assembled more deterministically, irrespective of core category, suggesting stronger host selection and/or more rapid dispersal during microbiome establishment. Core microbiota also generally showed stronger associations with postlarval survival than total communities or non-core taxa, although contrasting survival associations among core members indicated that core microbiota are not ecologically homogeneous. Our work provides the first systematic categorization and characterization of larval core microbiota throughout the shrimp larviculture cycle, offering new insights into the ecological significance of core microbiota and identifying potential universal targets for microbiome-based intervention in shrimp larviculture.

## 1. Introduction

Increasing attention has been directed toward the establishment of early-life microbiome and its potential long-term effects on host health across a broad range of species, especially in mammals (Malmuthuge et al., 2015; Yao et al., 2021). Previous studies have found that dysbiosis or imbalance of the intestinal microbial community in infants is associated with obesity (Houtman et al., 2022), chronic enteritis (Pammi et al., 2017) and immune system diseases (Fujimura et al., 2016; Gensollen et al., 2016). Meanwhile, early-life microbial interventions—such as vaginal microbiota transplantation and the modulation of microbial exposure through nutrition, probiotics/prebiotics, and prudent antibiotic use—can, to some extent, reshape the host microbiome and thus improve health (Davis et al., 2017; Dominguez-Bello et al., 2016). In recent years, the relationship between early-life microbiome establishment and the health of aquaculture animals has received increasing attention. For instance, a study has shown that an early-life core bacterial taxon (*Sulfitobacter* sp.) promote the growth of the sea cucumber (*Apostichopus japonicus*) (Yu et al., 2023); early microbial intervention has been found to shape the immune system of the Pacific oyster (*Crassostrea gigas*), thereby conferring lifelong and even transgenerational disease resistance (Fallet et al., 2022). Although the intestinal microbiota of Pacific white shrimp (*Penaeus vannamei*), one of the most productive aquaculture species worldwide (FAO, 2025), has long been a focus of research at the grow-out stage—particularly regarding its associations or causality with host health, growth, stress/disease resistance, and immunity (Zhang, Z. et al., 2025; Zhou et al., 2025)—comparatively little attention has been given to its early-life microbiota assembly throughout the larviculture cycle, which likely plays a critical role in determining the survival of postlarvae (as the product of larviculture) and consequently affects culture performance at the grow-out stage (Shan et al., 2025).

Currently, shrimp larviculture practices face a significant bottleneck in the poor reproducibility of larviculture batches with high final larval (postlarval) survival rates (Reyes et al., 2022). This issue may primarily stem from the following aspects: (1) the stochastic dynamics and divergence of microbiota and environmental conditions (Chen et al., 2025; Heyse et al., 2021); (2) the heavy reliance on practitioners’ experiential knowledge in managing and regulating larviculture systems, largely due to the lack of clear regulatory targets (eg. specific microbial taxa or guilds) and relevant guidance. Fundamentally, this stems from insufficient understanding of the larviculture microbial ecosystem, particularly the larval holobiont. Previous studies have demonstrated fine-scale succession and divergence patterns, assembly mechanisms, and major sources of the larval bacterial community of *P. vannamei* (Chen et al., 2025; Wang et al., 2020; Zhang, F. et al., 2025), while other studies have examined the associations between bacterial community and larval survival or diseases (Reyes et al., 2022; Shan et al., 2025); the assembly and sources of bacteria in the rearing water in the larviculture system (Heyse et al., 2021); and how probiotic or disinfectant treatments affect water and/or larval microbial community structure (Duan et al., 2020; Wang et al., 2020). However, a global understanding of the dynamics of larval microbial communities across developmental stages and geographic locations remains lacking. This gap hinders the establishment of a consensus on the key components of microecological regulation in shrimp larviculture systems.

Manipulating intestinal microbiota to enhance shrimp growth and to improve resistance to stresses/pathogens has been proposed and tested as a promising regulatory strategy for achieving microbiota homeostasis during the grow-out stage of culture (Dong et al., 2023; Guo et al., 2023). However, this approach remains largely unexplored in shrimp larviculture stage, though the microbial community composition alone can predict a more remarkable fraction of the variation in larval survival rates, compared with that during the grow-out stage (Shan et al., 2025). This limitation is primarily due to the lack of clearly defined and characterized microbial targets in the larviculture stage. With the rapid development of microbial ecology driven by high-throughput sequencing and multi-omics, the concept of “core microbiome” has evolved from its initial use in identifying the most common microbial members within a specific host population to becoming one of the central paradigms in host-microbe interaction research (Neu et al., 2021; Risely, 2020). In terms of taxonomic units, three mainstream definitions—regarding three categories of core microbiota—include common core, temporal core, and ecological core, respectively considering high occupancy, persistence, and network centrality of the sets of taxa within the microbiota of a given host population (Risely, 2020). Despite the empiricism in the defining parameters for these categories of core microbiota (Custer et al., 2023), these core taxa are hypothesized to play crucial functional roles in host physiology, metabolism, and immunity, thereby influencing host health and consequently emerging as attractive targets for microbiota-based manipulation (Zhang and Sun, 2022). For example, synthetic microbial communities (SynCom) constructed based on core or health-associated taxa have been applied to promote growth and/or to enhance resistance to pathogens/stresses of a range of hosts, including mammals (eg. mice, (Brugiroux et al., 2017)), terrestrial plants (watermelon, (Qiao et al., 2024)), aquatic plants (eg. macroalgae, (Wang et al., 2024)), and aquatic animals (eg. shrimp, (Guo et al., 2023)). However, our understanding remains limited regarding the presence, composition, dynamics, and assembly processes of the major categories of core members in the microbiota of *P. vannamei* larvae across different developmental stages and geographic regions. Importantly, the relationships and distinctions among different categories of core microbiota defined under varying criteria—and their associations with larval survival—are still poorly understood. These limitations are largely due to the lack of comprehensive datasets spanning major producing regions worldwide.

In this study, we integrated publicly available 16S rRNA gene amplicon datasets with additional self-generated datasets, originated from Asia (primarily China, accounting for 98.76% and 100% of larval and water samples) and Latin America (Ecuador). We aimed to test three hypotheses: (1) despite variation in prokaryotic community composition across geographic regions and larval developmental stages, shrimp larvae harbor a relatively conserved set of dominant taxa; (2) different categories of core taxa are distinct in their taxonomic composition, diversity, developmental-stage specificity, and ecological roles; and (3) compared with non-core taxa, core taxa are assembled more deterministically, exhibit stronger associations with postlarval survival, and therefore could play a more important role in larval microbiome assembly and host performance.

## 2. Materials and methods

### 2.1 Literature screening and 16S rRNA gene sequencing data retrieval

Publicly available 16S rRNA gene sequencing data on *P. vannamei* larviculture were retrieved by searching relevant literature published before 12 November 2024 in the Web of Science (WoS) database (https://www.webofscience.com/), following the screening procedure shown in Fig. S1. Specifically, the ‘Topic’ and ‘Title’ fields of the WoS search engine were queried using boolean combinations of terms related to research field (microbiome*, microbiota, bacterial community, microbial community, bacterial communities, microbial communities), target species (shrimp, *Penaeus vannamei*, *Litopenaeus vannamei*), and aquaculture stage (larva*, larviculture). We then screened the hit articles to select *P. vannamei* larviculture studies with (1) a clear statement to distinguish larval developmental stages (eg. *nauplius*, *zoea*, *mysis*, or *postlarvae*) and (2) publicly accessible 16S rRNA gene amplicon data of larvae and/or rearing water generated by Illumina sequencing platforms. Eventually, a total of 13 16S rRNA gene datasets covering variable regions that include the V4 region were selected. These comprised 11 datasets from peer-reviewed publications (including one published by our laboratory after the WoS search, referred to as Study 11) and two previously unpublished datasets generated by our laboratory, referred to as Study 12 and Study 13 (Table S1). Detailed information of these two additional larviculture batches and 16S rRNA gene sequencing data generation are provided in the Supplementary methods [see Supplementary information file].

### 2.2 Sample information curation and sequence processing

Key information of the datasets used for the meta-analysis were curated manually and then summarized in Tables S1 and S2. Samples with ambiguous information of developmental stages were excluded from the final dataset for meta-analysis. All raw 16S rRNA gene sequencing data were reassigned with customized IDs prior to processing. The dataset with paired-end reads were merged with the script *fastq_mergepairs* of USEARCH v11.0.667. To retrieve as many sequences as possible from different datasets while ensuring fair comparisons across datasets, e-PCR was performed to extract the V4 region of the 16S rRNA gene with the universal primers 515FY (5’-GTGYCAGCMGCCGCGGTAA-3’) and 806RB (5’-GGACTACNVGGGTWTCTAAT-3’) (Apprill et al., 2015; Parada et al., 2016) using the USEARCH script *search_pcr2* allowing two mismatches. When the raw sequences targeting the V3-V4 or V4 region contained only a single-end primer sequence, Cutadapt (Martin, 2011) or the USEARCH script *fastx_truncate* was used to ensure precise trimming for the single-end primer sequence. For all the datasets, sequences with lengths outside the range of 150-350 bp after e-PCR or trimming were removed using SeqKit v1.3-1 (Shen et al., 2016), and sequence recovery rates of the datasets were summarized in Table S2.

The sequences obtained after the above procedures from all datasets were pooled and subsequently processed using USEARCH v11.0.667. Quality filtering of the sequences was performed with a maximum expected error threshold of 1.0, followed by dereplication, using the script *fastx_uniques*. Denoising, chimera removal, and Zero-radius Operational Taxonomic Units (ZOTUs) identification were conducted with the script *usearch11-unoise3* (minsize = 4) based on the UNOISE3 algorithm (Edgar, 2016). All the sequences were mapped to ZOTUs at 100% identity with the script *usearch11 -otutab* to generate the ZOTU abundance table. Subsequently, The ZOTU sequences were aligned using the script *usearch11 -super5*, and then a phylogenetic tree was constructed by FastTree (Price et al., 2010). Taxonomic assignment of ZOTUs was performed using the QIIME 2 feature-classifier plugin (classify-sklearn), implementing a scikit-learn naïve Bayes classifier (Bokulich et al., 2018) trained on the Ribosomal Database Project (RDP) 16S rRNA reference training set (version 19) (Wang and Cole, 2024). The ZOTUs identified as Eukaryote, Mitochondria, and Chloroplast, as well as those taxonomically unassigned, were removed. The final prokaryotic ZOTU table comprised 888 samples (469 larval samples, 419 rearing water samples), containing 32,344 ZOTUs and 43,180,056 qualified reads (ranging 2637-181,678 reads per sample, mean = 48,626 reads per sample). The ZOTU table was rarefied at 8135 reads per sample for alpha-diversity analyses, and 8 water samples from Study 1 in Wenzhou and 1 water sample from Study 6 in Shenzhen with shallow sequencing depth (2637-7503) were excluded. Alpha-diversity indices, including ZOTU richness and Shannon-Wiener index, were calculated using the R package ‘vegan’ (Oksanen et al., 2025). Other analyses were conducted using total sum scaling (TSS) normalization as recommended (McKnight et al., 2019), unless otherwise indicated.

### 2.3 Identification of the common, temporal, and ecological core taxa in the larval prokaryotic community

The core ZOTUs of larval prokaryotic communities were identified regarding the definitions of three categories: common core, temporal core, and ecological core (Risely, 2020). Common core ZOTUs for the samples from China or other countries at pre-mouth opening stage (*nauplius*) and three post-mouth opening stages (*zoea*, *mysis*, and *postlarvae*) were identified based on the occupancy of ZOTUs (≥ 80%) throughout the samples of a given stage using the R package ‘tidyverse’ (Wickham et al., 2019). Venn diagrams were then created to show core ZOTUs for the samples from China or other countries at each stage, and the core ZOTUs shared by China and other countries were defined as the common core set for a given stage. The combination of the common core sets regarding the four major developmental stages was considered as the pan-common core set. The occupancy threshold for detecting core ZOTUs was determined based on a widely used definition of the common core microbiome as previously reported (Neu et al., 2021), while considering trade-offs between prevalence and the potential influence of outliers among samples. To further test the robustness of the core ZOTU thresholds, number of pan-common core ZOTUs and their proportional distribution among different taxonomic groups in larval samples, number of common core ZOTUs across stages, and relative abundance of taxonomic groups within the pan-common core sets were compared across occupancy thresholds ranging from 50% to 100%. The temporal core ZOTUs were defined as the persistent set of common core ZOTUs across the three post-mouth opening stages.

Ecological core ZOTUs are defined as taxa exerting disproportionate effects on microbial community (aka. potential keystone taxa) (Banerjee et al., 2018), whose removal or introduction may induce cascading changes in community diversity and structure (Dominguez-Garcia et al., 2019). Based on this definition, ecological core taxa can be identified using network centrality metrics associated with connectivity and information transfer efficiency (Agler et al., 2016). For each developmental stage, the ZOTUs with an occupancy greater than 20% across samples were retained for network inference to reduce the suspicious correlations. Pairwise correlations among ZOTUs were calculated using FastSpar (Watts et al., 2019), and only sound and significant associations (|r| > 0.4, *P* ≤ 0.001) were retained to construct the co-occurrence networks. Centrality metrics of network nodes (ZOTUs), including degree centrality, betweenness centrality, and closeness centrality, were calculated using Cytoscape v3.8.1. Consensus nodes falls into the upper 10% of a normal distribution of all the three metrics were identified as ecological core ZOTUs, representing taxa with high topological importance (Agler et al., 2016). The role of these ZOTUs in network stability was tested by comparing the decreasing extent of natural connectivity of the network after the sequential removal of ecological core or random nodes using the R package ‘igraph’ (Csárdi et al., 2026), and the significance of difference between the two node-removal procedures were assessed with Analysis of Covariance (ANCOVA) using the R package ‘stats’.

Venn diagrams were created to illustrate the overlaps and distinctions between common and ecological core ZOTUs. Heatmaps were generated to visualize the temporal persistence or stage specificity of the common core ZOTUs (including temporal core ZOTUs) and ecological core ZOTUs, their associations with postlarval survival rate (as described below), and their relative abundances across developmental stages and geographic locations, using the R package ‘ComplexHeatmap’ (Gu, 2022).

### 2.4 Inference of assembly processes of the core taxa into the larval prokaryotic community

The Sloan neutral modelling with the rarefied ZOTU tables as mentioned above was applied to infer the assembly processes of common (including temporal) and ecological core ZOTUs from different sources into the larval prokaryotic community across developmental stages. Specifically, the larva-to-larva model was used to infer processes among larvae, with the larval prokaryotic metacommunity as the source community. In contrast, the water-to-larva model was used to infer processes from rearing water to larvae, with the water prokaryotic metacommunity as the source community. Model fitting was conducted using the R packages ‘minpack.lm’ and ‘Hmisc’ (Elzhov et al., 2023; Harrell, 2023). Bar plots were used to show the proportion of ZOTU numbers regarding ecological process types and family-level taxonomic composition among common core, ecological core, and non-core ZOTUs using the R packages ‘dplyr’ (Wickham et al., 2023).

### 2.5 Fractionation of the core taxa according to their associations with postlarval survival rate

A multivariable linear model (MaAsLin2) was applied to assess the associations between individual ZOTUs and postlarval survival rate using the R package ‘Maaslin2’ (Mallick et al., 2021), with ZOTU abundance data subjected to TSS normalization and log transformation. Larviculture batches and geographic locations were included as covariates to account for potential confounding effects. Correlations were corrected with the Benjamini-Hochberg procedure, and ZOTUs with an adjusted *q* value < 0.05 were considered significantly correlated with postlarval survival rate. Volcano plot was then generated to fractionate pan-common core ZOTUs and pan-ecological core ZOTUs according to their associations with postlarval survival rate using the R package ‘ggplot2’.

The relative importance of the total prokaryotic community (represented by PC1 and PC2 scores derived from the Principal Coordinates Analysis (PCoA) plot) and different categories of core ZOTUs—including pan-common core set, pan-ecological core set, stage-specific common core sets, temporal core set, stage-specific ecological core sets, and non-core set—on postlarval survival rate was assessed using Random Forest regression implemented in the R package ‘randomForest’ (Liaw and Wiener, 2002). For each of the above sets, the ZOTUs that were significantly associated with postlarval survival rate were further fractionated and subjected to another Random Forest regression, allowing comparison of model performance before and after feature selection as well as shift in variable importance ranks. To improve the robustness of model evaluation under small-sample conditions, leave-one-out cross-validation combined with 10 repeated modeling runs was performed with ‘ntree = 1000’ and ‘mtry = 5’, and the mean of the 10 repeated predictions was used as the final predicted value.

### 2.6 General statistical analysis

All statistical analyses and data visualization were performed in the R environment (v4.4.2) unless otherwise indicated. The detailed procedures of general statistical analyses are provided in the Supplementary methods [see Supplementary information file].

## 3. Results and discussion

### 3.1 Sample distribution and attributes

This study collected publicly available 16S rRNA gene amplicon sequencing data from *P. vannamei* larviculture systems (including larval and rearing water samples) from the major producing regions worldwide, and integrated them with our own previously unpublished data. A total of 13 datasets meeting the screening criteria were obtained, comprising 888 samples (469 larval samples and 419 water samples; Fig. S1). The larval samples mainly originated from Wenzhou (Zhejiang Province), Shenzhen and Zhanjiang (Guangdong Province), and Wenchang and Sanya (Hainan Province), China (accounting for 84.65% in total), with additional contributions from Viluppuram, India and Guayaquil, Ecuador (Fig. 1). In contrast, the rearing water samples all came from China. The samples’ origins reflect China’s role as the world’s largest producer of *P. vannamei* (FAO, 2025). A total of 71 larval samples (15.14%) were diagnosed with diseases, including acute hepatopancreatic necrosis disease (AHPND, 47 samples) and Zoea-II syndrome (24 samples). Of 12 larviculture batches, 7 ones (involving 315 larval samples and 309 water samples) recorded the final larval (postlarval) survival rate.

**Figure 1.**
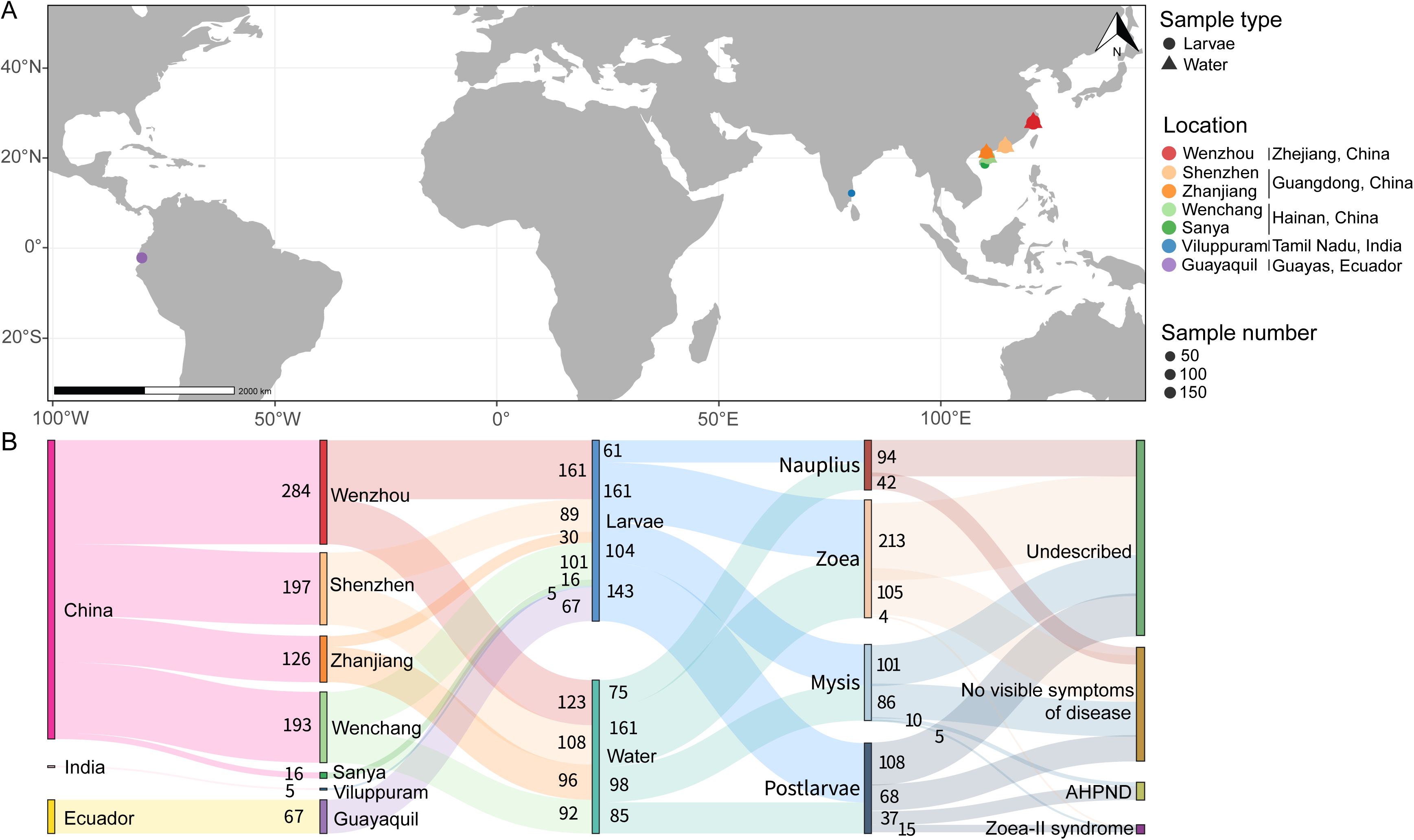
Geographic distribution and attributes of microbiota samples regarding *Penaeus vannamei* larviculture for the meta-analysis. Geographic distribution of shrimp larval and rearing water samples (A). Sankey diagram shows the flow of samples through different attributes, including country, city, sample type, developmental stage, health status (B). Width of the flow indicates sample size, the links between nodes represent the distribution of samples across different sample attributes. AHPND, acute hepatopancreatic necrosis disease.

### 3.2 Inconsistent developmental trends of larval prokaryotic alpha diversity across geographic locations

The Analysis of Variance (ANOVA) results showed that geographic location had the greatest effect on prokaryotic alpha-diversity indices (ZOTU richness and Shannon-Wiener index) in larvae and rearing water, followed by developmental stage and their interaction (all *P* < 0.001; Table S3). In contrast, sample type (larva vs. water) had a relatively weak effect on ZOTU richness, but a stronger influence on the Shannon-Wiener index than geographic location and developmental stage (based on *F* values, Table S3). A previous study (Study 3 in Table S1) reported a U-shaped pattern in larval bacterial alpha diversity across the complete developmental cycle, characterized by higher diversity at early (*nauplius*) and late (*postlarvae*) stages but lower levels during the intermediate stages (Wang et al., 2020). This trend paralleled the dynamics in alpha diversity observed in the rearing water and could be attributed to the community restructuring after the mouth opening of larvae. However, in this meta-analysis, inconsistent trends of larval prokaryotic alpha diversity over the developmental cycle were observed across geographic locations (Shenzhen, Zhanjiang, and Sanya) (Fig. S2). In addition, different larviculture batches from the same location (Wenzhou or Shenzhen) often exhibited distinct levels of alpha diversity at a given developmental stage (Fig. S3). These differences prompt us to examine how larval microbial diversity relates to that of the rearing water. Among the paired larva-water samples (265 pairs, primarily from Wenzhou, Shenzhen, and Wenchang), no significant correlation was observed between larval and water prokaryotic alpha diversity when samples across the entire developmental cycle were considered (Fig. 2). When analyzed on a stage-specific basis, associations between larval and water prokaryotic alpha diversity were barely observed after larvae opened their mouths, with the exception of a weak positive association at the postlarval stage for the Shannon-Wiener index (R^2^ = 0.120, *P* = 0.006; Fig. 2). This overall decoupling of prokaryotic alpha diversity between larvae and water partially explains the geographic heterogeneity in developmental trends of larval prokaryotic alpha diversity. Water environmental conditions were found associated with larval bacterial alpha diversity; however, most of the variance in alpha diversity remained unexplained (Chen et al., 2025). Collectively, the key determinants shaping larval prokaryotic alpha diversity during host development remain largely unrevealed and warrant further investigation, particularly with respect to the complex interplays among host genetics, larviculture management practices (e.g., feeding regimes and system maintenance), water environmental conditions, and the water microbiota.

**Figure 2.**
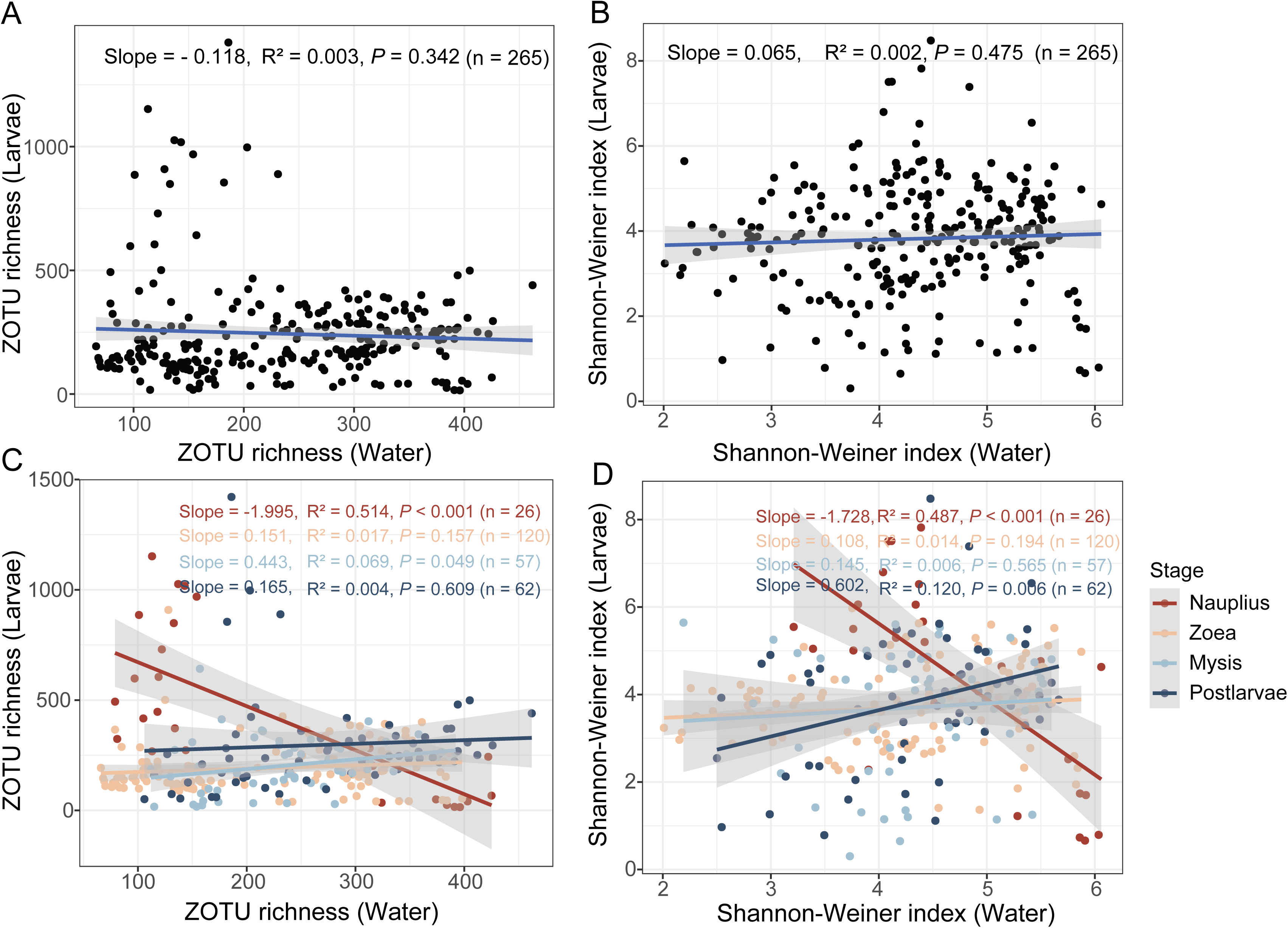
Relationships of prokaryotic alpha diversity between paired larval and rearing water samples across developmental stages. The linear regression analyses illustrate the overall relationships between prokaryotic alpha diversity in larvae and their rearing water (A and B), as well as those at different larval developmental stages (C and D). The shaded areas represent the 95% confidence intervals.

### 3.3 Compositional variation of larval prokaryotic communities across developmental stages and geographic locations

The larval prokaryotic communities were dominated by bacteria, accounting for 99.90% in the entire dataset (Fig. 3). At the high taxonomic level, *Alphaproteobacteria*, *Gammaproteobacteria*, and *Bacteroidota* ranked as the three most abundant bacterial groups in larvae across nearly all geographic locations. An exception was observed in larval samples (all zoeal samples) from Viluppuram, India, where the three most abundant bacterial groups were *Gammaproteobacteria*, *Bacillota*, and *Bacteroidota*. *Bacillota* was occasionally dominant in Shenzhen and Zhanjiang (mainly during the *zoea*, *mysis*, and *postlarvae* stages) and in Wenchang (during the *nauplius* and *postlarvae* stages), China, as well as in Viluppuram, India (at the *zoea* stage). Overall, *Roseobacteraceae* (mean relative abundance: 37.71%), *Vibrionaceae* (10.27%), and *Flavobacteriaceae* (9.63%) were the three most abundant bacterial families in larvae across developmental stages and geographic locations, together accounting for 57.62% of the entire larval dataset (Fig. 3). The taxonomic composition at the family level exhibited pronounced geographic and developmental variation (Fig. 3 and Table S4). Therefore, larval communities across most locations displayed distinct yet consistently *Roseobacteraceae*-dominated compositions, with the exception of those from Viluppuram, India (Fig. 3). In China, *Roseobacteraceae* was the most dominant family across all surveyed locations, including Wenzhou (mean relative abundance: 29.88%), Shenzhen (30.53%), Zhanjiang (43.53%), Wenchang (56.81%), and Sanya (39.16%). Other prevalent families—such as *Flavobacteriaceae*, *Vibrionaceae*, *Pseudoalteromonadaceae*, *Moraxellaceae*, and *Weeksellaceae*—varied in relative abundance among locations. Outside China, larval samples from Guayaquil, Ecuador, were similarly dominated by *Roseobacteraceae* (36.64%), followed by *Vibrionaceae* (16.69%) and *Flavobacteriaceae* (4.77%). In contrast, larval communities from Viluppuram, India, exhibited a distinct composition, characterized by the predominance of *Moraxellaceae* (36.66%) and *Caryophanaceae* (27.60%), with *Roseobacteraceae* relatively less contributing 6.78% of the total community. Similarly, a conserved set of dominant bacterial families prevailed in the rearing water (Find more details in Supplementary Results and Fig.S4).

**Figure 3.**
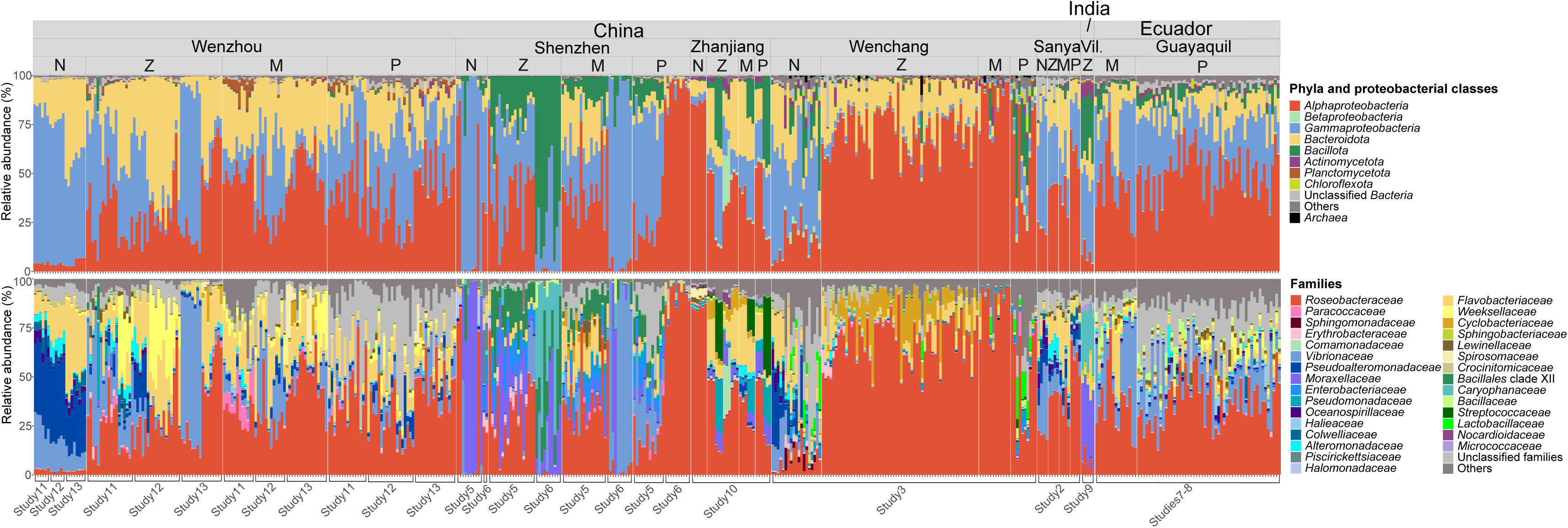
Taxonomic distribution of larval prokaryotic communities across geographic locations and developmental stages. The upper and bottom panels show phylum- and family-level taxonomic composition, respectively, with *Proteobacteria* (now *Pseudomonadota*) further classified at the class level. Taxa with average relative abundance ≥ 2% at least at one developmental stage in one city are shown. N, *nauplius*; Z, *zoea*; M, *mysis*; P, *postlarvae*; Vil., Viluppuram.

Prokaryotic community composition (based on ZOTUs) in both larvae and rearing water varied significantly with geographic location, larval developmental stage, sample type, and their interactions, regardless of beta-diversity metrics (Fig. 4 and Table 1). The consistent compositional variation in larval microbial communities over the developmental cycle suggests an inherent development-associated succession of larval microbiota. Our meta-analysis firstly reveals that geographic location exerted a stronger influence compared with developmental stage regardless of sample sets (R^2^_PERMANOVA_ = 0.236 vs. 0.126, 0.201 vs. 0.094, and 0.127 vs. 0.018 for larval, water, and all samples, all *P* = 0.001, based on Bray-Curtis dissimilarity; Table 1). Among all the larval samples, significant distance-decay patterns in taxonomic and phylogenetic similarities of prokaryotic communities between geographic locations were detected at each developmental stage, with the strength of this pattern decreasing as host development progressed (from *nauplius* and *zoea* to *mysis* and *postlarvae*; Fig. S5). In addition, distance-decay patterns in larval and water communities were also sound when exclusively considering Chinese samples.

**Figure 4.**
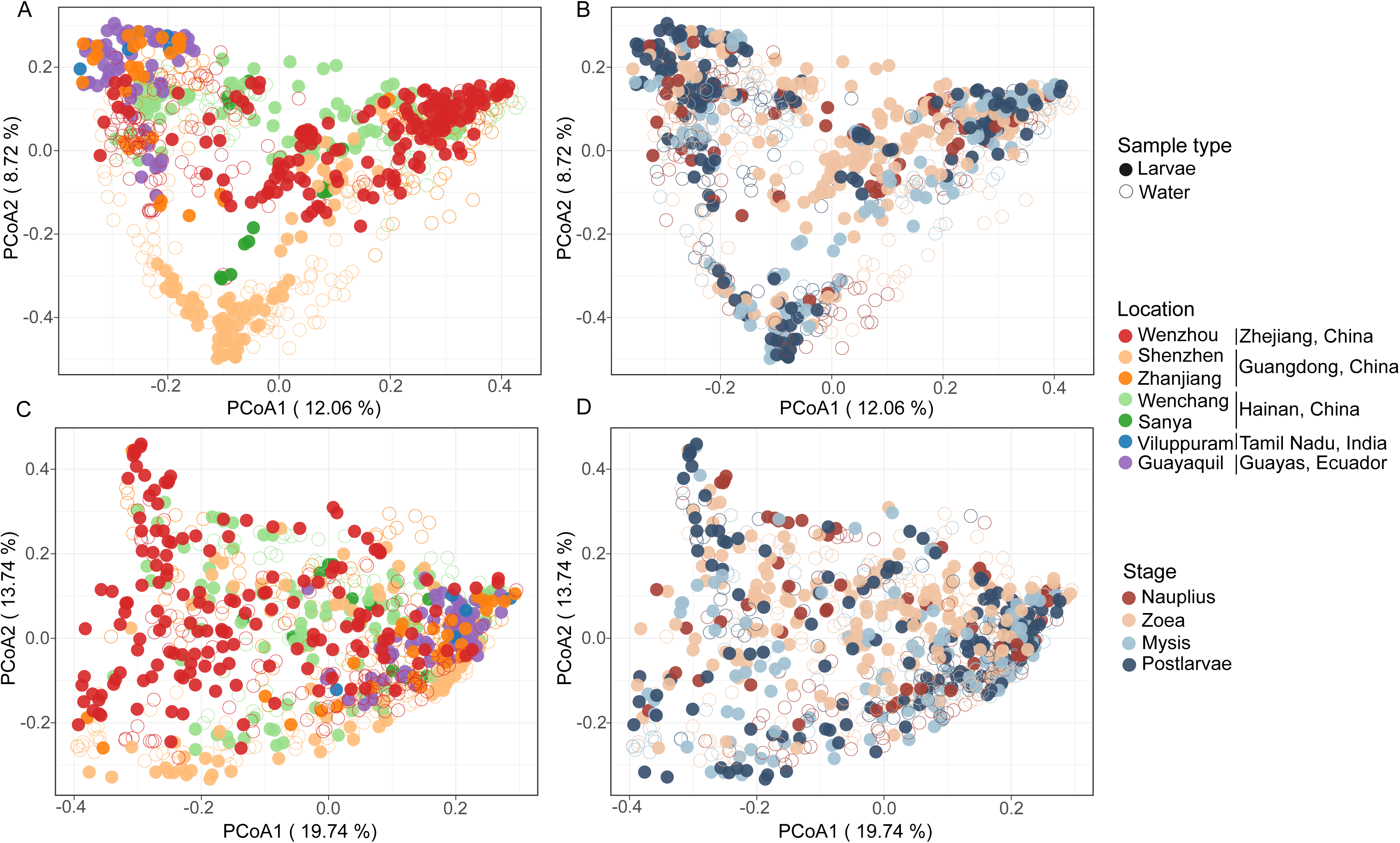
Prokaryotic community variation in larvae and rearing water across geographic locations and developmental stages. Principal Coordinate Analysis (PCoA) based on Bray-Curtis dissimilarity (A and B) and weighted UniFrac distance (C and D) illustrating compositional variation of larval and water prokaryotic communities across geographic locations (left panels) and developmental stages (right panels).

**Table 1.**
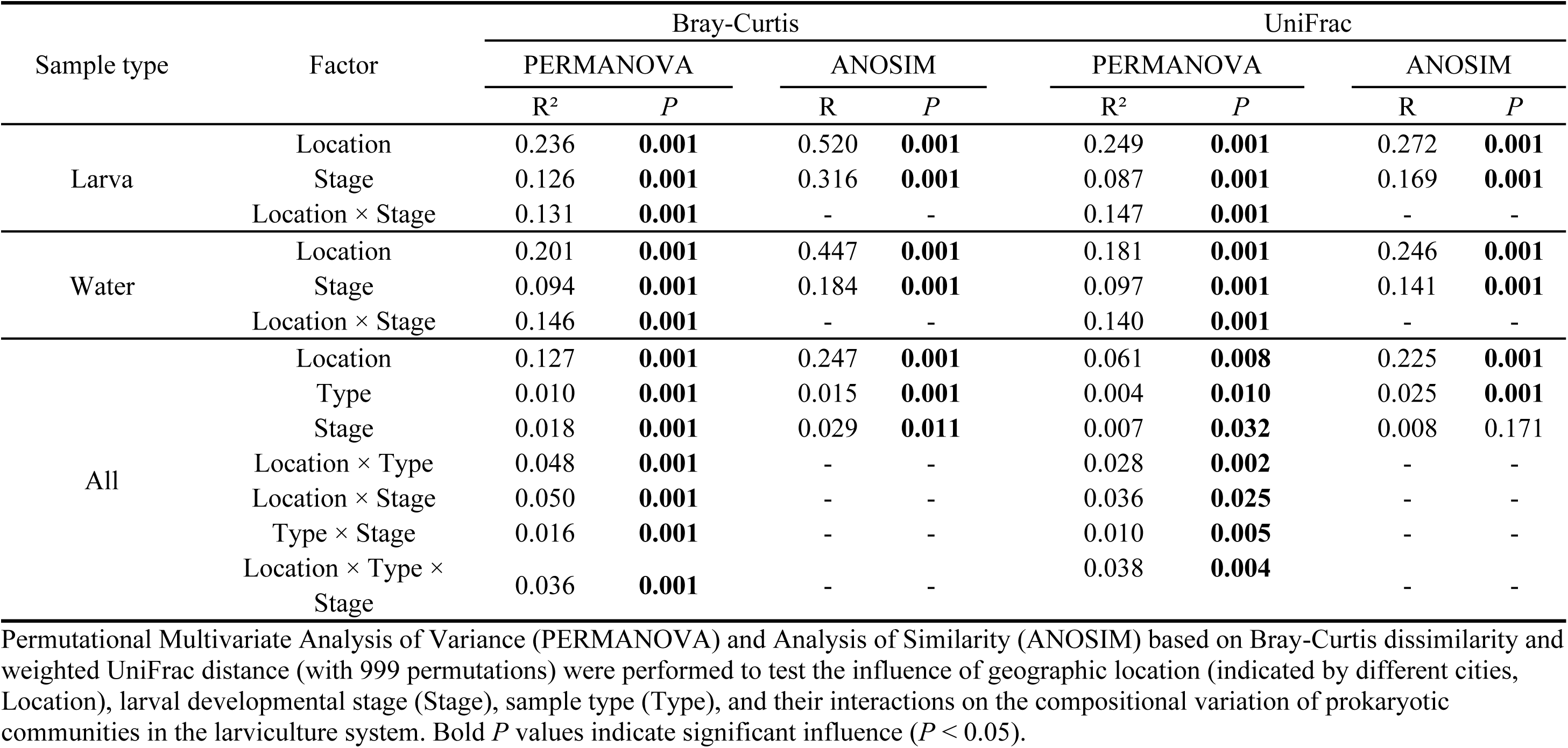
Effects of three major factors on prokaryotic community compositions in the larviculture system.

Larval bacterial community composition has been reported to be shaped by the combined and interactive effects of developmental stage, water environmental conditions, and water microbiota (Chen et al., 2025). Therefore, differences in environmental conditions and microbiota in rearing water among geographic locations may be associated with the observed biogeographic patterns. However, water environmental profiles were not comprehensively characterized or well overlapped among most datasets, which limited further analysis. On the other hand, although water samples were limited to those from China, larval and rearing water prokaryotic communities shared several common taxonomic features, particularly at higher taxonomic ranks (Supplementary Results, Fig. S4). Both habitats were dominated by a limited number of bacterial lineages, with *Alphaproteobacteria*, *Gammaproteobacteria*, and *Bacteroidota* consistently representing major components of the communities. At the family level, *Roseobacteraceae* and *Flavobacteriaceae* were prevalent in both larvae and water, suggesting that these bacterial groups constitute a conserved microbial pool within larviculture systems and may serve as important sources for larval microbial colonization (Zhang, F. et al., 2025). Despite these shared features, pronounced differences in community composition were evident between larvae and water (Figs. 3 and S4; Table S4). Combined with the earlier point that larval alpha diversity trends were largely decoupled from those in water, this shows that the larval microbiota is not a simple reflection of the ambient microbial environment.

### 3.4 Stage specificity and temporal persistency of larval common core taxa

Given the dynamic succession of larval microbiota with host development, it is essential to identify core taxa and examine their specificity and persistence across developmental stages. On this basis, we further propose the concept of a pan-common core set—the combination of common core taxa regarding different stages—to represent the common core microbial assemblage over the entire larviculture cycle. When identifying common core microbiota, the ubiquity of specific taxa across a set of communities were often assessed using occupancy as an indicator (Neu et al., 2021; Risely, 2020). However, the choice of occupancy threshold is largely empirical, often ranging from 50% to 100% (Bragina et al., 2015; Keady et al., 2021; Pfeiffer et al., 2017). To balance a stricter standard with necessary flexibility (due to the presence of outlier samples), we adopted an 80% threshold for identifying common core taxa. We also tested performance across thresholds from 50% to 100% and found that when the threshold reached up to 90% and 100%, no core taxa were detected for certain developmental stages (Fig. S6A), supporting the justification of the 80% threshold in a way.

For larval prokaryotic communities, 24, 7, 22, and 26 common core ZOTUs were identified for the pre-mouth opening stage (*nauplius*, for which only Chinese samples were available) and three post-mouth opening stages (*zoea*, *mysis*, and *postlarvae*), respectively, yielding a total of 35 pan-common core ZOTUs (Fig. 5A). Overall, the pan-common core ZOTUs exhibited broad ubiquity across different stages: 19 ZOTUs were identified as core members in three or four stages, whereas 16 ZOTUs showed specificity to only one or two stages. After the mouth opening of larvae, the common core set of Chinese samples generally represented the core set of all samples at *mysis* and *postlarvae* stages, while only 7 of the 24 core ZOTUs of Chinese samples overlapped with the overall core set at *zoea* stage. Previous studies have identified the *zoea* stage as a critical tipping point in both the composition and sources of larval bacterial communities as well as a key window for the colonization of external bacteria (Wang et al., 2020; Zhang, F. et al., 2025), which together may explain the reduced number of common core taxa observed at this stage compared with others stages. Furthermore, 6 out of 7 ZOTUs in the zoeal common core set were also identified as common core taxa in the other two post-mouth opening stages, and thus the zoeal core set can largely define the temporal core set over host development.

**Figure 5.**
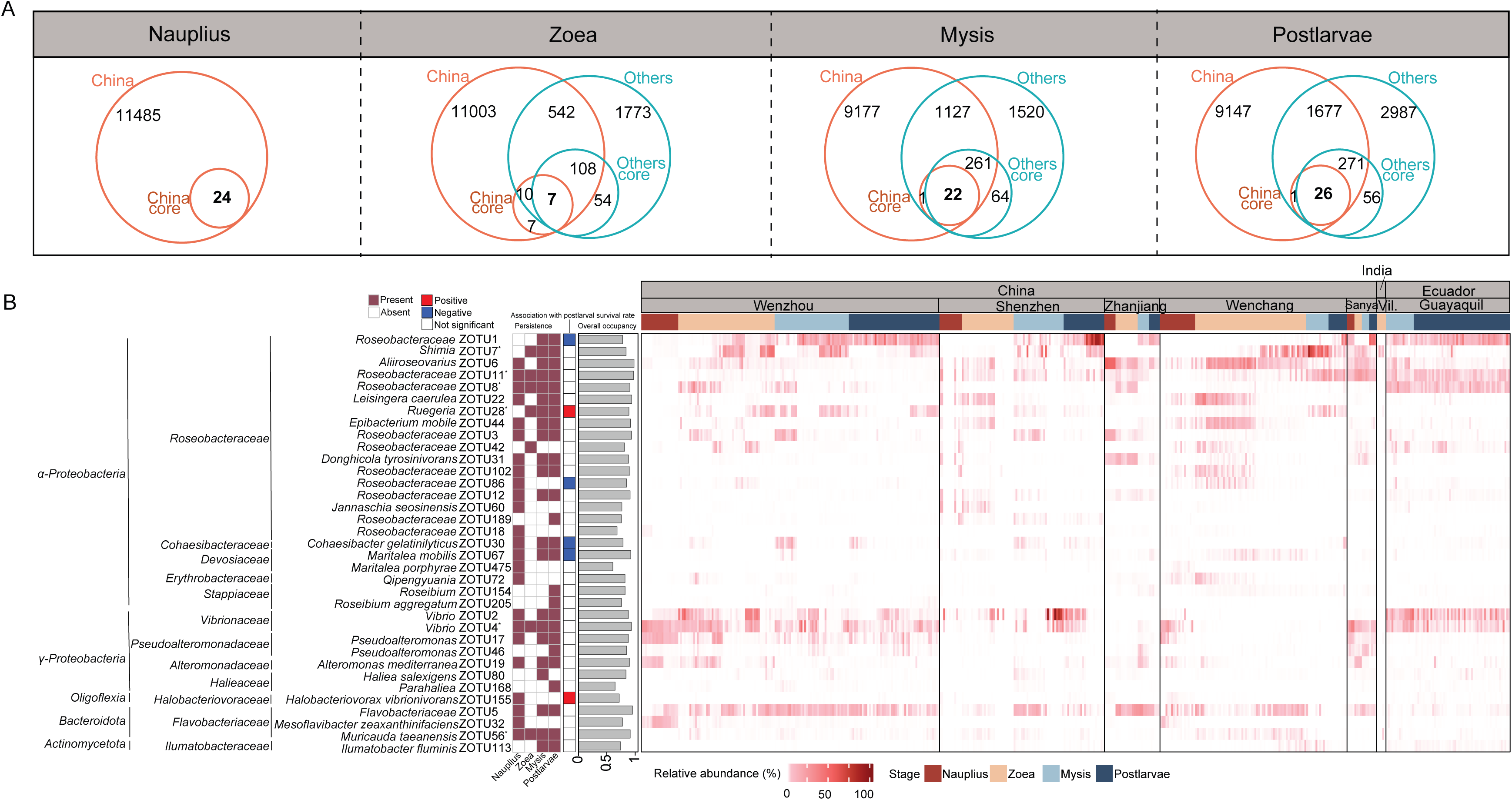
Identification of common and temporal core prokaryotic ZOTUs in larvae across four major developmental stages. The Venn diagrams in panel (A) show ZOTUs with an occupancy ≥ 80% at each stage for the samples from China or other countries, with bolded values indicating the common core ZOTUs shared by China and other countries. The left heatmap in panel (B) illustrates the presence (dark rose cells) or absence (blank cells) of 35 pan-common core ZOTUs (the combined set of common core ZOTUs of four stages) in the core set at a given stage; the ZOTU IDs with asterisk represent temporal core ZOTUs consistently included in the common core sets of three post-mouth opening stages; the middle heatmap illustrates the associations between the ZOTUs and postlarval survival rate (For the complete results of the correlation analysis, see Fig. S8 in the Supplementary information); the bar plots shows overall occupancy of the ZOTUs across all larval samples; and the right heatmap illustrates the relative abundance of pan-common core ZOTUs in larval prokaryotic communities across geographical locations and host developmental stages.

### 3.5 *Roseobacteraceae* dominated the diverse common core microbiota of shrimp larvae

Regardless of the occupancy threshold used to identify “common core taxa” (50%-100%), *Roseobacteraceae* taxa consistently dominated the larval pan-common core microbiota, in terms of average proportion within the pan-common core set (21.47-65%) and average relative abundance in the prokaryotic community (13.86-31.04%) or the pan-common core set (45.02-69.14%) (Fig. S6B-C). At the 80% threshold, 17 of 35 larval pan-common core ZOTUs were affiliated with *Roseobacteraceae*, including members of the genera *Shimia*, *Aliiroseovarius*, *Leisingera* (*L. caerulea*), *Ruegeria*, *Epibacterium* (*E. mobile*), *Donghicola* (*D. tyrosinivorans*), and *Jannaschia* (*J. seosinensis*), with one ZOTU assigned to each genus, as well as 10 ZOTUs that could not be resolved at the genus level (Fig. 5). These *Roseobacteraceae* ZOTUs accounted for 55.29% and 25.06% of cumulative relative abundance in the pan-common core set and the prokaryotic community. In addition, four of the six temporal core ZOTUs were affiliated with *Roseobacteraceae* (Fig. 5). Members of *Roseobacteraceae* (aka. the *Roseobacter* group, formerly classified within *Rhodobacteraceae* (Liang et al., 2021)) are frequently detected at high abundance in rearing water and shrimp-associated microbiota at both larviculture and grow-out stages (Dong et al., 2023; Wang et al., 2020). Accumulating evidence indicates that *Roseobacteraceae* taxa can contribute to host health through pathogen inhibition (Sonnenschein et al., 2021) and metabolic support (Buchan et al., 2026). Several roseobacters, including *Phaeobacter* sp. and *Ruegeria* sp., are capable of producing antibacterial compounds such as tropodithietic acid (TDA), which can suppress the growth of pathogenic *Vibrio* species and thus improve the survival of shrimp and fish (Dittmann et al., 2020; Sonnenschein et al., 2017). These properties have led to increasing interest in *Roseobacteraceae* as promising aquaculture probiotic candidates (Sonnenschein et al., 2021). In addition, many *Roseobacteraceae* members possess strong capacities for organic matter transformation and vitamin biosynthesis, and actively participate in carbon and sulfur cycling in aquaculture systems (Rajeev et al., 2024). Such metabolic versatility suggests that *Roseobacteraceae* may also facilitate nutrient utilization and other host-associated metabolic processes in shrimp (Sonnenschein et al., 2021). However, not all members of *Roseobacteraceae* are beneficial. Certain taxa, such as *Aliiroseovarius crassostreae*, the causative agent of Roseovarius oyster disease (ROS), have been identified as pathogens in aquatic animals (Kessner et al., 2016), highlighting substantial functional differentiation within the family during host-associated lifestyles. These findings indicate that finer taxonomic and functional resolution of *Roseobacteraceae* is necessary to distinguish beneficial symbionts from potentially pathogenic members. As a family of well-studied marine heterotrophy (Buchan et al., 2026), their specific functions in shrimp larvae remain poorly understood. Such high taxonomic diversity and abundance of *Roseobacteraceae* ZOTUs in the larval common and temporal core microbiota suggest their crucial and diversified functions in shrimp larvae. Further studies are therefore needed to clarify functional contributions and host interactions of distinct *Roseobacteraceae* lineages during shrimp larval development.

Besides *Roseobacteraceae*, the pan-common core microbiota of shrimp larvae (Fig. 5) included bacterial taxa with potentially beneficial, pathogenic, and yet-unclear host-associated roles. *Pseudoalteromonas* (ZOTUs 17 and 46) is well known for producing antimicrobial compounds and extracellular enzymes that inhibit pathogenic *Vibrio* spp. and support host digestion and immune defense (Gustilatov et al., 2024; Khanjani et al., 2024). *Halobacteriovorax vibrionivorans* (ZOTU155) is a predatory bacterium capable of preying on Gram-negative bacteria, including *Vibrio*, and may therefore contribute to pathogen control in aquaculture systems (Ye et al., 2019). *Alteromonas mediterranea* (ZOTU19) has been associated with organic matter degradation and secretion of bioactive metabolites (Zhang and Sun, 2025), while *Mesoflavibacter zeaxanthinifaciens* (ZOTU32) and *Muricauda taeanensis* (ZOTU56) may additionally contribute to antioxidant production through carotenoid biosynthesis (Asker et al., 2007; Endo et al., 2023). In contrast, *Vibrio* (ZOTUs 2 and 4) contains well-known opportunistic pathogens frequently related to shrimp diseases and microbiota dysbiosis, while *Maritalea* (ZOTUs 67 and 475) has been reported to increase in the intestines of shrimp when resisting white spot syndrome (Wanna et al., 2025), suggesting a possible association with disease-related microbial shifts. Other taxa, including *Haliea salexigens* (ZOTU80), *Parahaliea* (ZOTU168), *Qipengyuania* (ZOTU72), *Ilumatobacter fluminis* (ZOTU113), *Cohaesibacter gelatinilyticus* (ZOTU30), and *Roseibium* (ZOTUs 154 and 205) remain poorly characterized in shrimp-associated microbiota. Overall, these findings highlight the high taxonomic and phylogenetic diversities and potential functional differentiation of larval common core microbiota and the need for further studies on their host-associated functions.

### 3.6 Ecological core taxa as potential keystones in the microbiota of shrimp larvae

A total of 64 pan-ecological core ZOTUs were identified based on the consensus of three network centrality metrics (Agler et al., 2016), including 21, 16, 13, and 22 for the *nauplius*, *zoea*, *mysis,* and *postlarvae* stages, respectively (Fig. 6A). The importance of these ZOTUs in network stability were confirmed by the dramatically enhanced decreasing trend of network natural connectivity (indicating robustness of the network to perturbation (Dominguez-Garcia et al., 2019)) after gradually removing nodes regarding these ZOTUs compared with random node removals (Fig 6B), suggesting their potential as keystones in microbiota stability (Wang et al., 2023). As defined differently, overlaps between ecological and common core ZOTUs were rarely found at each stage, except for 2 and 1 ZOTU(s) at *nauplius* and *zoea* stages, respectively (Fig. S7). Overall, 8 ZOTUs (2 *Roseobacteraceae* ZOTUs: *Leisingera caerulea* and unclassified genus; 2 *Devosiaceae* ZOTUs: *Maritalea mobilis* and *M. porphyrae*; 1 *Roseibium* ZOTU; 2 *Vibrio* ZOTUs; and 1 *Mesoflavibacter zeaxanthinifaciens* ZOTU) were included in both pan-common core and pan-ecological core sets. The pan-ecological core ZOTUs showed generally lower occupancy (40.29% in average) and cumulative relative abundance (0.63% in average; Fig 6C) compared with those of the pan-common core set (84.20% and 1.30%; Fig 5B). These results suggest that ecological core taxa are not necessarily characterized by high occupancy or dominance, but instead align with the key hypothesis of keystone taxa that they exert disproportionately important ecological effects relative to their abundance (Banerjee et al., 2018).

**Figure 6.**
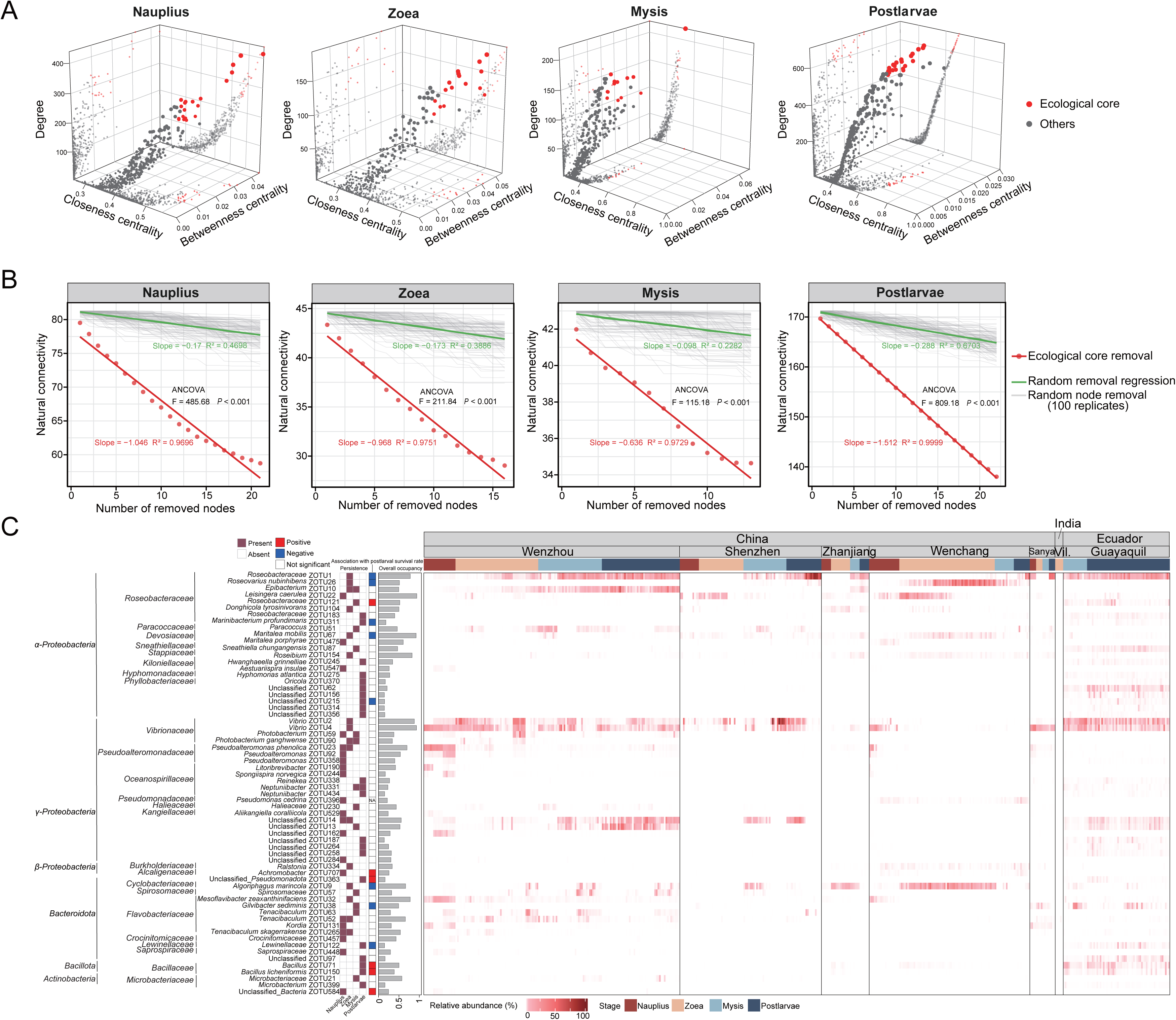
Identification of ecological core ZOTUs in larval prokaryotic communities across four major developmental stages. Three-dimensional plots distinguish ecological core and non-ecological core ZOTUs based network centrality metrics, including degree centrality, betweenness centrality, and closeness centrality for each developmental stage (A). The role of ecological core ZOTUs on network invulnerability was tested by comparing the decreasing extent of natural connectivity of the network after the sequential removal of ecological core or random nodes (with 100 replicates) (B). The significantly steeper linear regression (with larger absolute slope) as tested by Analysis of Covariance (ANCOVA) indicates dominating influence of ecological core ZOTUs on network invulnerability. The left heatmap in panel (C) illustrates the presence (dark rose cells) or absence (blank cells) of 64 pan-ecological core ZOTUs (the combined set of ecological core ZOTUs of four stages) in the core set at a given stage; the middle heatmap illustrates the associations between the ZOTUs and postlarval survival rate (For the complete results of the correlation analysis, see Fig. S8 in the Supplementary information); the bar plots shows overall occupancy of the ZOTUs across all larval samples; and the right heatmap illustrates the relative abundance of pan-ecological core ZOTUs in larval prokaryotic communities across geographical locations and host developmental stages.

The ecological core ZOTUs exhibited even higher phylogenetic diversity and stronger stage specificity than the common core microbiota. These ZOTUs were distributed across a wide range of bacterial lineages, with particularly high representation of *Roseobacteraceae*, *Oceanospirillaceae*, *Vibrionaceae*, *Flavobacteriaceae*, and *Pseudoalteromonadaceae* (Fig. 6C). Among them, *Roseobacteraceae* included members affiliated with *Marinibacterium*, *Roseovarius*, *Leisingera*, *Donghicola*, *Epibacterium*, and several genus-unresolved taxa, highlighting the ecological importance and functional diversification of roseobacter during shrimp larval development. Other ecological core members representatively included *Pseudoalteromonas*, *Neptuniibacter*, *Vibrio*, *Photobacterium*, *Tenacibaculum*, *Bacillus*, and *Maritalea*. Notably, 15 ecological core ZOTUs could not be classified at the family or higher taxonomic ranks, suggesting that shrimp larvae harbor a considerable proportion of phylogenetically novel or yet-uncultivated bacterial lineages that may represent undescribed taxa. The composition of ecological core taxa changed markedly across developmental stages (Fig. 6C). The *nauplius* stage exhibited the most taxonomically dispersed ecological core set, dominated by *Pseudoalteromonas*, *Tenacibaculum*, and *Oceanospirillaceae* taxa, together with multiple low-frequency lineages from distinct families. During the *zoea* stage, *Roseobacteraceae* became one of the dominant ecological core groups, accompanied by persistent *Tenacibaculum*, *Pseudoalteromonas*, and *Vibrionaceae* taxa. In the *mysis* stage, the ecological core set became relatively simplified, mainly consisting of *Roseobacteraceae*, *Photobacterium*, and *Flavobacteriaceae* taxa. In contrast, the *postlarvae* stage showed increased representation of *Neptuniibacter*, *Bacillus*, and taxa (10 ZOTUs) unclassified at higher taxonomic ranks. Despite the strong succession and continuity of the larval microbiota with host development (Zhang, F. et al., 2025), only a few ecological core ZOTUs persisted across adjacent developmental stages, indicating that microbial interaction networks may undergo repeated restructuring during larval development and molting.

Functionally, the ecological core taxa potentially included both beneficial and harmful microorganisms. Several taxa, including *Pseudoalteromonas*, roseobacter-related genera (*Leisingera*, *Roseovarius*, and *Epibacterium*), and *Bacillus*, have previously been associated with antimicrobial activity, nutrient cycling, and vitamin biosynthesis in marine animal (De Santiago et al., 2026; Kesarcodi-Watson et al., 2008; Ringø, 2020), suggesting potential probiotic or symbiotic roles during larval development. In contrast, members of *Vibrio*, *Photobacterium*, and *Tenacibaculum* include known opportunistic pathogens or disease-associated taxa in aquatic organisms (Echeverria-Bugueno et al., 2023; Mabrok et al., 2023). This observation supports the idea that pathogens may also exhibit keystone traits if they tend to restructuring a healthy microbiota into a dysbiotic state (Hajishengallis et al., 2012). Therefore, ecological core or keystone properties should not be interpreted exclusively as indicators of beneficial functions, but may also reflect taxa with pathogenic or destabilizing ecological effects.

Importantly, the pronounced turnover and stage specificity of ecological core ZOTUs suggest that identifying temporally persistent keystone taxa in shrimp larvae may require alternative approaches beyond static co-occurrence network analyses, such as time-series ecological modeling, interaction-based inference, and experimental perturbation studies (Amit and Bashan, 2023; Banerjee et al., 2018). Furthermore, whether the ecological core ZOTUs identified here truly function as keystone taxa remains to be experimentally validated. Isolation and cultivation of representative strains, particularly the numerous unclassified taxa detected in this study, together with controlled colonization and functional assays, will be essential for determining their actual ecological roles and impacts on shrimp larval health and microbiome stability.

### 3.7 Assembly processes of core taxa into the microbiota of shrimp larvae

Evaluating the assembly processes of core taxa among shrimp larvae and from rearing water into larval microbiota may help reveal host selection preferences toward specific core taxa, as well as differences in the dispersal capacities of core taxa from different sources. In this study, regardless of developmental stage, the overall larval prokaryotic communities generally fit the neutral model when larval microbiota were used as the source community (larva-to-larva model, Fig. 7A-7D). In contrast, both the common core and ecological core taxa were largely governed by non-neutral processes, as substantial proportions of ZOTUs occurred both above and below model predictions, whereas non-core taxa were predominantly neutrally distributed. These results suggest that, relative to non-core taxa, the incorporation of core taxa into larval microbiota may involve stronger deterministic processes, potentially reflecting host-mediated selection or exclusion. On the other hand, because our meta-analysis includes samples from geographically distinct hatcheries, these patterns may also partly reflect differences in dispersal ability—leading to different degrees of dispersal limitation—among core taxa during transmission between larvae. Interestingly, even closely related core taxa, particularly within *Roseobacteraceae*, distributed among distinct ecological sub-divisions and were presumed to be governed by different ecological processes, indicating potential differences in host preference and/or dispersal capability. Such ecological differentiation may correspond to the functional diversification discussed above.

**Figure 7.**
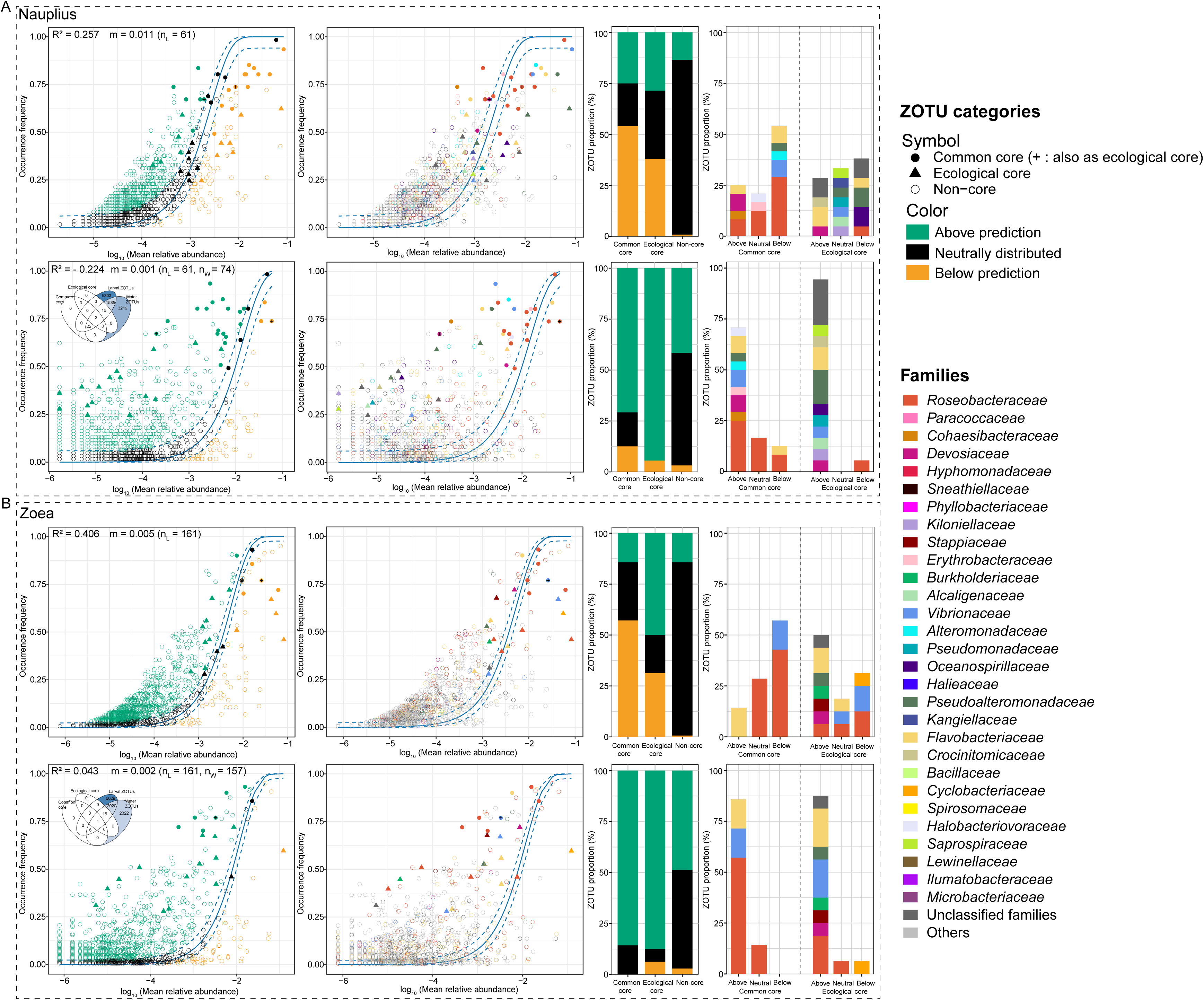
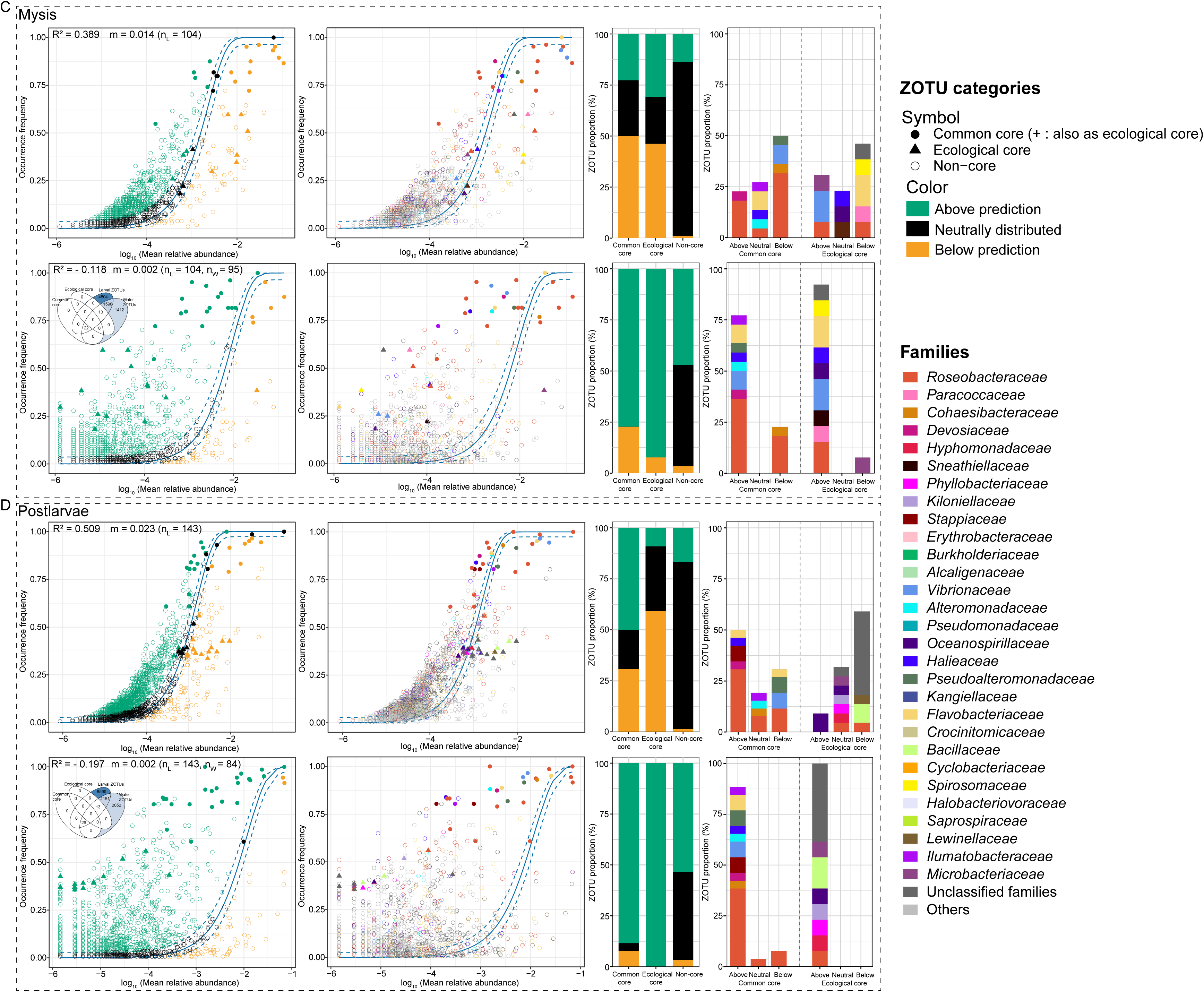
Neutral model inference of larval prokaryotic community assembly processes across developmental stages and source metacommunities. The models were fitted using either the larval metacommunity (larva-to-larva model, top plots) or the water metacommunity (water-to-larva model, bottom plots) as the source community across the *nauplius* (A), *zoea* (B), *mysis* (C), and *postlarvae* (D) stages. The R² value indicates the goodness of fit of the neutral model, ranging from 0 (no fit) to 1 (perfect fit). The m value indicates the migration rate of taxa into the larval prokaryotic community from the source community. n_L_ and n_W_ indicate the numbers of larval and water samples used for model fitting, respectively. The Venn diagrams show the overlap among the common core, ecological core, larval, and water ZOTUs, and the ZOTUs shared by water and larval samples were included in the fitting of water-to-larva models. Dashed lines denote the 95% confidence intervals of the model predictions, and ZOTUs falling within these confidence intervals are considered to be neutrally distributed. ZOTUs occurring more frequently than predicted by the model are shown in green, while those occurring less frequently than predicted are shown in orange. Stacked bar plots show the ZOTU proportions of common core ZOTUs, ecological core ZOTUs, and non-core ZOTUs across the three model-defined subsets, as well as the three subsets of common core ZOTUs and ecological core ZOTUs classified at the family level. Circles with black cross denote the ZOTUs included in both common core and ecological core sets for a given stage.

When neutral models were constructed using water microbiota as the source community (water-to-larva model), microbial assembly from water into larval microbiota generally deviated from neutral expectations across the four major developmental stages (Fig. 7). Generally, a considerable proportion of water-derived ZOTUs appeared to be deterministically assembled (non-neutral vs. neutral, *nauplius*: 45.78% vs. 54.22%; *zoea*: 52.11% vs. 47.89%; *mysis*: 51.56% vs. 48.44%; *postlarvae*: 57.40% vs. 42.60%), and both common core and ecological core taxa were predominantly enriched above neutral predictions (70.83-88.46% and 87.5-100%, respectively), suggesting stronger host filtering (selection) and/or more rapid dispersal of most core taxa in(to) larvae. Notably, some core taxa, particularly two *Vibrio* ZOTUs (ZOTUs 2 and 4), showed opposite patterns between the larva-to-larva and water-to-larva models, being underrepresented in larval transmission but overrepresented from water to larvae (Fig. 7A-7D). This pattern may indicate that these taxa are less efficiently transmitted among larvae or excluded by host, but more readily transmitted (arguably in a passive way) from the surrounding environment. This finding suggests that the control of potentially pathogenic core taxa may need to begin with management of the rearing water microbiota. In addition, a number of ecological core taxa, particularly during the *nauplius* and *postlarvae* stages, were not detected in rearing water, suggesting that some core microbes may originate from alternative sources beyond the surrounding water microbiota.

### 3.8 Associations between core microbiota and postlarval survival rate

Since the concept of the core microbiota was proposed, its ecological roles and functional significance in host health have remained controversial and lack broad consensus (Neu et al., 2021; Risely, 2020). This uncertainty has also limited the practical application of core microbiota as potential targets for microbiome manipulation to improve host health. In particular, it remains unclear whether core taxa consistently contribute to host fitness, merely reflect stable environmental colonizers, or include conditionally pathogenic members with context-dependent effects (Sanders-Smith et al., 2020; Zhou et al., 2024). As discussed above, both potentially beneficial and harmful bacterial taxa have been identified in both common and ecological core sets. Here, we further evaluated the relationships between core microbiota and shrimp larval performance by examining both individual core taxa and different core sets as integrated microbial assemblages in relation to postlarval survival rate. Such analyses may help distinguish potentially beneficial, detrimental, and functionally neutral members within the core microbiota, thereby providing a more comprehensive understanding of how core taxa contribute to larval health. Furthermore, identifying core microbiota subsets that are strongly associated with postlarval survival may help pinpoint key microbial taxa and provide a foundation for the targeted design of simplified synthetic microbial consortia aimed at stabilizing microbiome homeostasis in shrimp larviculture systems.

The combined pan-common core and pan-ecological core sets comprised 91 ZOTUs, of which 18 were significantly correlated with postlarval survival rate, including 8 positively correlated and 10 negatively correlated ZOTUs (Figs 5, 6, and S8). Among the positively correlative ZOTUs, six belonged to the ecological core set, including taxa affiliated with *Roseobacteraceae*, *Bacillus* (including *B. licheniformis*), *Achromobacter*, and unclassified members of *Pseudomonadota* and even *Bacteria* (Fig. 6 and S8). In addition, one positively correlative ZOTU belonged to the common-temporal core taxon *Ruegeria*, while another one corresponded to the *nauplius*-stage common core taxon *Halobacteriovorax vibrionivorans*. Among the negatively correlative ZOTUs, six belonged to the ecological core set, including *Gilvibacter sediminis*, *Algoriphagus marincola*, *Roseovarius nubinhibens*, *Marinibacterium profundimaris*, and unclassified members of *Lewinellaceae* and *Hyphomicrobiales*. Two negatively correlative ZOTUs belonged to the common core set, including *Cohaesibacter gelatinilyticus* and one *Roseobacteraceae* taxon, while two additional taxa, including *Maritalea mobilis* and another *Roseobacteraceae* taxon, were shared between the common core and ecological core sets.

Random Forest analysis revealed that, among the different categories of core microbiota, the ecological core sets of the *nauplius*, *zoea*, and *postlarvae* stages, together with the zoeal common core set (including all temporal core taxa), showed relatively more important on postlarval survival rate than other core sets, the total prokaryotic community, or the non-core set when evaluated as integrated microbial assemblages (Fig. 8A). This finding suggests that specific subsets of the microbiota may contribute disproportionately to larval performance, whereas a substantial proportion of the overall community may be functionally redundant or only weakly associated with survival-related variation. Further fractionation of each core set by retaining only survival-associated ZOTUs further increased their predictive importance (Fig. 8B). Notably, the fractionated zoeal common/temporal core subset, which consisted solely of a single *Ruegeria* ZOTU (survival-positive), together with the mysis and postlarval ecological core subsets (including survival-positive taxa affiliated with *Roseobacteraceae*, *Bacillus*, and unclassified *Pseudomonadota* and survival-negative taxa affiliated with *Gilvibacter sediminis*, *Marinibacterium profundimaris*, and unclassified members of *Lewinellaceae* and *Hyphomicrobiales*), exhibited the strongest associations with postlarval survival. These results highlight the importance of refining core microbiota into smaller subsets that are more closely linked to host performance, rather than treating the entire core assemblage as ecologically homogeneous (Risely, 2020; Yin et al., 2025). The stage-specific shifts in influential core subsets also suggest that different core taxa may become important at different stages, potentially reflecting dynamic restructuring of microbial interactions over larval development. Although the present study did not directly assess microbial functions, these patterns raise the possibility that temporal core and ecological core taxa may act in a relay-like manner over larval development, thereby contributing to microbiome stability across developmental transitions. Future studies integrating cultivation, functional validation, and multi-omics approaches will be necessary to determine whether these survival-associated core taxa represent functionally important component of the shrimp larval microbiota. On the other hand, although abundance-based analyses revealed that only a small proportion of core taxa were associated with postlarval survival, the ecological importance of core microbiota, particularly the ecological core taxa, may not primarily depend on their abundance, but rather on their roles in mediating microbial interactions within the community. This aspect also deserves further investigation in future.

**Figure 8.**
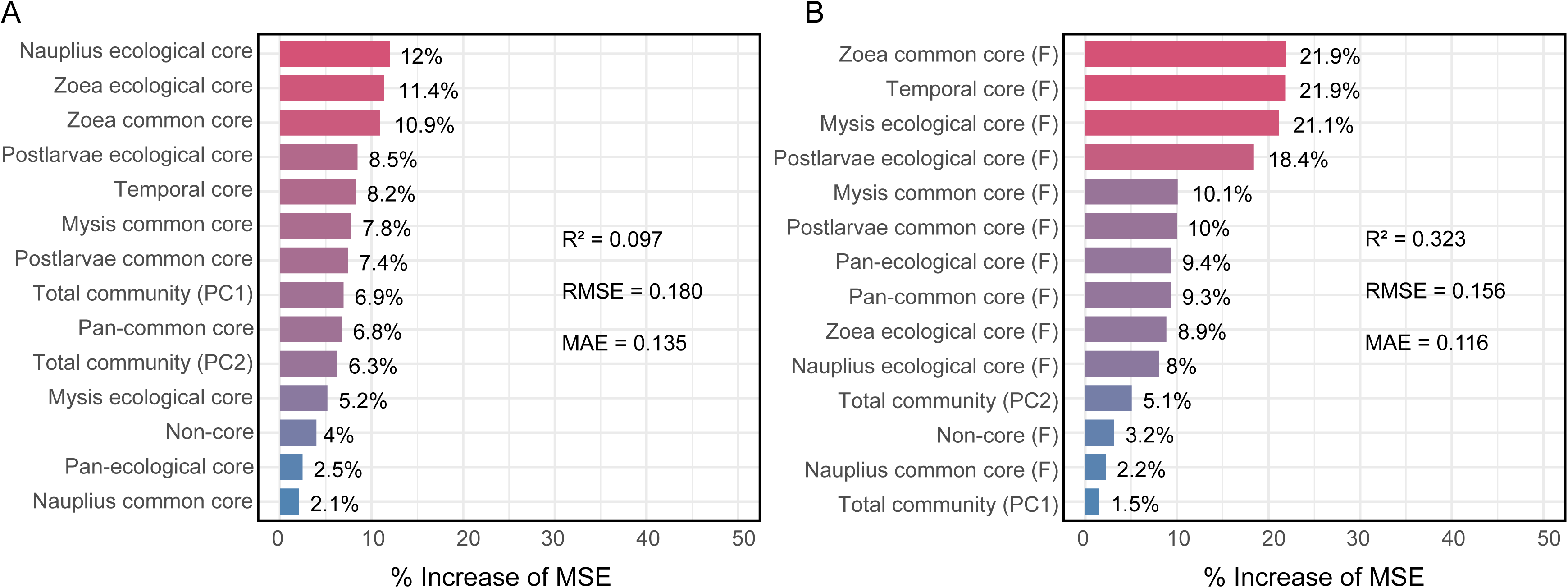
Associations of total prokaryotic community features and core ZOTU sets with shrimp postlarval survival. Random Forest Regression analysis illustrates the predictive relationships between the final larval (postlarval) survival rate and microbial features, including total prokaryotic community composition (indicated by the first two principal coordinate scores (PC1 and PC2) based on Bray-Curtis dissimilarity as shown in Fig.4) and the cumulative relative abundance of different categories of core ZOTUs in postlarval samples (n = 123) (A). For each category of core set, ZOTUs that are significantly associated with postlarval survival rate (as shown in Fig. S8) were further fractionated (F) and subjected to another Random Forest analysis (B). The percentage increase in MSE (Mean Square Error) for each variable represents the relative increase in the model’s MSE after permuting the values of that variable, and was used to assess the importance of the variable in predicting the outcome. R^2^ measures the proportion of variance explained by the model, with values ranging from 0 to 1, where higher values indicating better fit; RMSE (Root Mean Square Error) evaluates the standard deviation of prediction errors, where smaller values indicate higher accuracy; and MAE (Mean Absolute Error) calculates the average absolute prediction error, with smaller values reflecting better performance.

## Conclusions

This study provides the first systematic categorization and characterization of larval core microbiota throughout the larviculture cycle of *P. vannamei* at a global scale. Despite strong geographic variation in larval prokaryotic communities, a conserved set of dominant bacterial taxa consistently persisted across developmental stages and geographic locations. By integrating occupancy, persistence, and ecological network centrality, we demonstrated that common, temporal, and ecological core microbiota differ substantially in composition, diversity, stage specificity, and ecological roles in network stability. The core taxa were more strongly shaped by deterministic processes and showed closer associations with postlarval survival than the total community or non-core taxa, suggesting their potential importance for larval health and survival. However, the contrasting survival associations among different core members indicate that core microbiota should not be regarded as ecologically homogeneous, and that finer-scale fractionation is necessary to identify functionally relevant microbial targets. Overall, our findings advance the ecological understanding of larval core microbiota assembly in shrimp hatcheries and provide potential universal targets for microbiome-based intervention strategies to improve larval performance and production stability.

## Author contributions

K.W. conceived and strategized the meta-analysis. K.W., J.Y., C.C., D.Z., X.H., and M.Y. organized the additional larviculture monitoring; R.C., H.S., and Y.Z. collected the additional samples and did the bench work; R.C., W.Z., H.S., and K.W. analyzed the data; K.W., W.Z., and R.C. wrote the manuscript. All the authors reviewed and approved the manuscript.

## Ethical statement

This study is primarily a meta-analysis based on previously published data. Any shrimp larvae used to generate new data in our laboratory were handled in accordance with internationally recognized 3Rs principles (Replacement, Reduction, and Refinement), with every effort made to minimize animal use, stress, and suffering.

## Acknowledgements

Kai Wang dedicates this work in memory of his beloved golden retriever, Pangding (胖丁; April 29, 2018–January 8, 2026).

## Data availability

The accession numbers of all the public accessible sequence data are summarized in Table S2. Additional sequence data generated by this study have been submitted to the Sequence Read Archive of NCBI under accession number PRJNA1381294 and will be accessible with the following link after the acceptance of the manuscript: https://www.ncbi.nlm.nih.gov/sra/PRJNA1381294.

## References

Agler, M.T., Ruhe, J., Kroll, S., Morhenn, C., Kim, S.T., Weigel, D., Kemen, E.M., 2016. Microbial hub taxa link host and abiotic factors to plant microbiome variation. PLoS Biol. 14 (1), e1002352. 10.1371/journal.pbio.1002352.

Amit, G., Bashan, A., 2023. Top-down identification of keystone taxa in the microbiome. Nat. Commun. 14, 3951. 10.1038/s41467-023-39459-5.

Apprill, A., McNally, S., Parsons, R., Weber, L., 2015. Minor revision to V4 region SSU rRNA 806R gene primer greatly increases detection of SAR11 bacterioplankton. Aquat. Microb. Ecol. 75, 129–137. 10.3354/ame01753.

Asker, D., Beppu, T., Ueda, K., 2007. Mesoflavibacter zeaxanthinifaciens gen. nov., sp. nov., a novel zeaxanthin-producing marine bacterium of the family Flavobacteriaceae. Syst. Appl. Microbiol. 30 (4), 291–296. 10.1016/j.syapm.2006.12.003.

Banerjee, S., Schlaeppi, K., Van Der Heijden, M.G., 2018. Keystone taxa as drivers of microbiome structure and functioning. Nat. Rev. Microbiol. 16 (9), 567–576. 10.1038/s41579-018-0024-1.

Bokulich, N.A., Kaehler, B.D., Rideout, J.R., Dillon, M., Bolyen, E., Knight, R., Huttley, G.A., Gregory Caporaso, J., 2018. Optimizing taxonomic classification of marker-gene amplicon sequences with QIIME 2’s q2-feature-classifier plugin. Microbiome 6, 90. 10.1186/s40168-018-0470-z.

Bragina, A., Berg, C., Berg, G., 2015. The core microbiome bonds the Alpine bog vegetation to a transkingdom metacommunity. Mol. Ecol. 24 (18), 4795–4807. 10.1111/mec.13342.

Brugiroux, S., Beutler, M., Pfann, C., Garzetti, D., Ruscheweyh, H.-J., Ring, D., Diehl, M., Herp, S., Lötscher, Y., Hussain, S., 2017. Genome-guided design of a defined mouse microbiota that confers colonization resistance against *Salmonella enterica* serovar *Typhimurium*. Nat. Microbiol. 2, 16215. 10.1038/nmicrobiol.2016.215.

Buchan, A., Gonzalez, J.M., Moran, M.A., 2026. Discoveries with *Roseobacteraceae*: bacterial models for ocean heterotrophy. Ann. Rev. Mar. Sci. 18, 381–408. 10.1146/annurev-marine-050823-105708.

Chen, H., Zhang, F., Yu, J., Chen, R., Zhang, D., Chen, C., Wang, K., 2025. Divergence patterns of bacterial communities between larviculture systems of two *Penaeus vannamei* strains with distinct culture traits. Aquaculture 606, 742572. 10.1016/j.aquaculture.2025.742572.

Csárdi, G., Nepusz, T., Traag, V., Horvát, S., Zanini, F., Noom, D., Müller, K., Schoch, D., Salmon, M., 2026. igraph: Network Analysis and Visualization in R, version 2.3.3 [software]. https://CRAN.R-project.org/package=igraph.

Custer, G.F., Gans, M., van Diepen, L.T.A., Dini-Andreote, F., Buerkle, C.A., 2023. Comparative analysis of core microbiome assignments: implications for ecological synthesis. mSystems 8, e01066–22. 10.1128/msystems.01066-22.

Davis, E.C., Wang, M., Donovan, S.M., 2017. The role of early life nutrition in the establishment of gastrointestinal microbial composition and function. Gut Microbes 8 (2), 143–171. 10.1080/19490976.2016.1278104.

De Santiago, A., Barnes, S., Pereira, T.J., Marcellino Barros, M., Durden, L., Han, M.K., Thrash, J.C., Bik, H.M., 2026. *Pseudoalteromonas* is a symbiont of marine invertebrates that exhibits broad patterns of phylosymbiosis. ISME J. 20 (1), wrag091. 10.1093/ismejo/wrag091.

Dittmann, K.K., Rasmussen, B.B., Melchiorsen, J., Sonnenschein, E.C., Gram, L., Bentzon-Tilia, M., 2020. Changes in the microbiome of mariculture feed organisms after treatment with a potentially probiotic strain of *Phaeobacter inhibens*. Appl. Environ. Microbiol. 86, e00499–20. 10.1128/AEM.00499-20.

Dominguez-Bello, M.G., De Jesus-Laboy, K.M., Shen, N., Cox, L.M., Amir, A., Gonzalez, A., Bokulich, N.A., Song, S.J., Hoashi, M., Rivera-Vinas, J.I., Mendez, K., Knight, R., Clemente, J.C., 2016. Partial restoration of the microbiota of cesarean-born infants via vaginal microbial transfer. Nat. Med. 22, 250–253. 10.1038/nm.4039.

Dominguez-Garcia, V., Dakos, V., Kefi, S., 2019. Unveiling dimensions of stability in complex ecological networks. Proc. Natl. Acad. Sci. U. S. A. 116 (51), 25714–25720. 10.1073/pnas.1904470116.

Dong, P., Guo, H., Huang, L., Zhang, D., Wang, K., 2023. Glucose addition improves the culture performance of Pacific white shrimp by regulating the assembly of Rhodobacteraceae taxa in gut bacterial community. Aquaculture 567, 739254. 10.1016/j.aquaculture.2023.739254.

Duan, Y., Tang, Y., Huang, J., Zhang, J., Lin, H., Jiang, S., Wang, R., Wang, G., 2020. Changes in the microbial community of *Litopenaeus vannamei* larvae and rearing water during different growth stages after disinfection treatment of hatchery water. J. Microbiol. 58, 741–749. 10.1007/s12275-020-0053-0.

Echeverria-Bugueno, M., Irgang, R., Mancilla-Schulz, J., Avendano-Herrera, R., 2023. Healthy and infected Atlantic salmon (*Salmo salar*) skin-mucus response to *Tenacibaculum dicentrarchi* under in vitro conditions. Fish Shellfish Immunol. 136, 108747. 10.1016/j.fsi.2023.108747.

Edgar, R.C., 2016. UNOISE2: improved error-correction for Illumina 16S and ITS amplicon sequencing. bioRxiv [preprint], 081257. 10.1101/081257.

Elzhov, T.V., Mullen, K.M., Spiess, A.-N., Bolker, B., 2023. minpack.lm: R Interface to the Levenberg-Marquardt Nonlinear Least-Squares Algorithm Found in MINPACK, Plus Support for Bounds, version 1.2-4 [software]. https://CRAN.R-project.org/package=minpack.lm.

Endo, F., Motone, K., Kai, K., Kitakaze, T., Nishikawa, Y., Nishimura, Y., Kitamura, R., Takagi, T., Ito, M., Miura, N., 2023. Characterization of carotenoid-producing *Muricauda* sp. strains isolated from the reef-building coral *Galaxea fascicularis* and subtropical seawater. bioRxiv [preprint], 2023.12.12.571358. 10.1101/2023.12.12.571358.

Fallet, M., Montagnani, C., Petton, B., Dantan, L., de Lorgeril, J., Comarmond, S., Chaparro, C., Toulza, E., Boitard, S., Escoubas, J.M., Vergnes, A., Le Grand, J., Bulla, I., Gueguen, Y., Vidal-Dupiol, J., Grunau, C., Mitta, G., Cosseau, C., 2022. Early life microbial exposures shape the *Crassostrea gigas* immune system for lifelong and intergenerational disease protection. Microbiome 10, 85. 10.1186/s40168-022-01280-5.

FAO, 2025. FishStat: Global aquaculture production 1950-2023. (released on 31 March 2025). https://www.fao.org/fishery/en/statistics/software/fishstatj.

Fujimura, K.E., Sitarik, A.R., Havstad, S., Lin, D.L., Levan, S., Fadrosh, D., Panzer, A.R., LaMere, B., Rackaityte, E., Lukacs, N.W., Wegienka, G., Boushey, H.A., Ownby, D.R., Zoratti, E.M., Levin, A.M., Johnson, C.C., Lynch, S.V., 2016. Neonatal gut microbiota associates with childhood multisensitized atopy and T cell differentiation. Nat. Med. 22, 1187–1191. 10.1038/nm.4176.

Gensollen, T., Iyer, S.S., Kasper, D.L., Blumberg, R.S., 2016. How colonization by microbiota in early life shapes the immune system. Science 352 (6285), 539–544. 10.1126/science.aad9378.

Gu, Z., 2022. Complex heatmap visualization. iMeta 1 (3), e43. 10.1002/imt2.43.

Guo, H., Fu, X., He, J., Wang, R., Yan, M., Wang, J., Dong, P., Huang, L., Zhang, D., 2023. Gut bacterial consortium enriched in a biofloc system protects shrimp against *Vibrio parahaemolyticus* infection. Microbiome 11, 230. 10.1186/s40168-023-01663-2.

Gustilatov, M., Widanarni, W., Ekasari, J., Julyantoro, P.G.S., Waturangi, D.E., 2024. Biofloc system supplemented by *Pseudoalteromonas piscicida* 1Ub protects the Pacific white shrimp *Penaeus vannamei* from *Vibrio parahaemolyticus* infection. Aquac. Fish. 9 (6), 967–974. 10.1016/j.aaf.2023.05.003.

Hajishengallis, G., Darveau, R.P., Curtis, M.A., 2012. The keystone-pathogen hypothesis. Nat. Rev. Microbiol. 10, 717–725. 10.1038/nrmicro2873.

Harrell, F.E., Jr., 2023. Hmisc: Harrell Miscellaneous, version 5.1-1 [software]. https://CRAN.R-project.org/package=Hmisc.

Heyse, J., Props, R., Kongnuan, P., De Schryver, P., Rombaut, G., Defoirdt, T., Boon, N., 2021. Rearing water microbiomes in white leg shrimp (*Litopenaeus vannamei*) larviculture assemble stochastically and are influenced by the microbiomes of live feed products. Environ. Microbiol. 23 (1), 281–298. 10.1111/1462-2920.15310.

Houtman, T.A., Eckermann, H.A., Smidt, H., de Weerth, C., 2022. Gut microbiota and BMI throughout childhood: the role of firmicutes, bacteroidetes, and short-chain fatty acid producers. Sci. Rep. 12, 3140. 10.1038/s41598-022-07176-6.

Keady, M.M., Prado, N., Lim, H.C., Brown, J., Paris, S., Muletz-Wolz, C.R., 2021. Clinical health issues, reproductive hormones, and metabolic hormones associated with gut microbiome structure in African and Asian elephants. Anim. Microbiome 3, 85. 10.1186/s42523-021-00146-9.

Kesarcodi-Watson, A., Kaspar, H., Lategan, M.J., Gibson, L., 2008. Probiotics in aquaculture: the need, principles and mechanisms of action and screening processes. Aquaculture 274 (1), 1–14. 10.1016/j.aquaculture.2007.11.019.

Kessner, L., Spinard, E., Gomez-Chiarri, M., Rowley, D.C., Nelson, D.R., 2016. Draft genome sequence of *Aliiroseovarius crassostreae* CV919-312, the causative agent of *Roseovarius Oyster* disease (formerly juvenile oyster disease). Genome Announc. 4 (2), e00148–16. 10.1128/genomea.00148-16.

Khanjani, M.H., Mozanzadeh, M.T., Gisbert, E., Hoseinifar, S.H., 2024. Probiotics, prebiotics, and synbiotics in shrimp aquaculture: their effects on growth performance, immune responses, and gut microbiome. Aquac. Rep. 38, 102362. 10.1016/j.aqrep.2024.102362.

Liang, K.Y., Orata, F.D., Boucher, Y.F., Case, R.J., 2021. Roseobacters in a sea of poly-and paraphyly: whole genome-based taxonomy of the family *Rhodobacteraceae* and the proposal for the split of the “Roseobacter clade” into a novel family, Roseobacteraceae fam. nov. Front. Microbiol. 12, 683109. 10.3389/fmicb.2021.683109.

Liaw, A., Wiener, M., 2002. Classification and regression by randomForest. R News 2 (3), 18–22. https://journal.r-project.org/articles/RN-2002-022/.

Mabrok, M., Algammal, A.M., Sivaramasamy, E., Hetta, H.F., Atwah, B., Alghamdi, S., Fawzy, A., Avendaño-Herrera, R., Rodkhum, C., 2023. Tenacibaculosis caused by *Tenacibaculum maritimum*: updated knowledge of this marine bacterial fish pathogen. Front. Cell. Infect. Microbiol. 12, 1068000. 10.3389/fcimb.2022.1068000.

Mallick, H., Rahnavard, A., McIver, L.J., Ma, S., Zhang, Y., Nguyen, L.H., Tickle, T.L., Weingart, G., Ren, B., Schwager, E.H., Chatterjee, S., Thompson, K.N., Wilkinson, J.E., Subramanian, A., Lu, Y., Waldron, L., Paulson, J.N., Franzosa, E.A., Bravo, H.C., Huttenhower, C., 2021. Multivariable association discovery in population-scale meta-omics studies. PLoS Comput. Biol. 17 (11), e1009442. 10.1371/journal.pcbi.1009442.

Malmuthuge, N., Griebel, P.J., Guan le, L., 2015. The gut microbiome and its potential role in the development and function of newborn calf gastrointestinal tract. Front. Vet. Sci. 2, 36. 10.3389/fvets.2015.00036.

Martin, M., 2011. Cutadapt removes adapter sequences from high-throughput sequencing reads. EMBnet J. 17 (1), 10–12. 10.14806/ej.17.1.200.

McKnight, D.T., Huerlimann, R., Bower, D.S., Schwarzkopf, L., Alford, R.A., Zenger, K.R., 2019. Methods for normalizing microbiome data: an ecological perspective. Methods Ecol. Evol. 10 (3), 389–400. 10.1111/2041-210X.13115.

Neu, A.T., Allen, E.E., Roy, K., 2021. Defining and quantifying the core microbiome: challenges and prospects. Proc. Natl. Acad. Sci. U. S. A. 118 (51), e2104429118. 10.1073/pnas.2104429118.

Oksanen, J., Simpson, G., Blanchet, F., Kindt, R., Legendre, P., Minchin, P., O’Hara, R., Solymos, P., Stevens, M., Szoecs, E., Wagner, H., Barbour, M., Bedward, M., Bolker, B., Borcard, D., Borman, T., Carvalho, G., Chirico, M., De Caceres, M., Durand, S., Evangelista, H., FitzJohn, R., Friendly, M., Furneaux, B., Hannigan, G., Hill, M., Lahti, L., Martino, C., McGlinn, D., Ouellette, M., Ribeiro Cunha, E., Smith, T., Stier, A., Ter Braak, C., Weedon, J., 2025. vegan: Community Ecology Package, version 2.7-2 [software]. https://CRAN.R-project.org/package=vegan.

Pammi, M., Cope, J., Tarr, P.I., Warner, B.B., Morrow, A.L., Mai, V., Gregory, K.E., Kroll, J.S., McMurtry, V., Ferris, M.J., Engstrand, L., Lilja, H.E., Hollister, E.B., Versalovic, J., Neu, J., 2017. Intestinal dysbiosis in preterm infants preceding necrotizing enterocolitis: a systematic review and meta-analysis. Microbiome 5, 31. 10.1186/s40168-017-0248-8.

Parada, A.E., Needham, D.M., Fuhrman, J.A., 2016. Every base matters: assessing small subunit rRNA primers for marine microbiomes with mock communities, time series and global field samples. Environ. Microbiol. 18 (5), 1403–1414. 10.1111/1462-2920.13023.

Pfeiffer, S., Mitter, B., Oswald, A., Schloter-Hai, B., Schloter, M., Declerck, S., Sessitsch, A., 2017. Rhizosphere microbiomes of potato cultivated in the High Andes show stable and dynamic core microbiomes with different responses to plant development. FEMS Microbiol. Ecol. 93 (2), fiw242. 10.1093/femsec/fiw242.

Price, M.N., Dehal, P.S., Arkin, A.P., 2010. FastTree 2--approximately maximum-likelihood trees for large alignments. PLoS One 5 (3), e9490. 10.1371/journal.pone.0009490.

Qiao, Y., Wang, Z., Sun, H., Guo, H., Song, Y., Zhang, H., Ruan, Y., Xu, Q., Huang, Q., Shen, Q., Ling, N., 2024. Synthetic community derived from grafted watermelon rhizosphere provides protection for ungrafted watermelon against *Fusarium oxysporum* via microbial synergistic effects. Microbiome 12, 101. 10.1186/s40168-024-01814-z.

Rajeev, M., Jung, I., Kang, I., Cho, J.-C., 2024. Genome-centric metagenomics provides insights into the core microbial community and functional profiles of biofloc aquaculture. mSystems 9 (10), e00782–24. 10.1128/msystems.00782-24.

Reyes, G., Betancourt, I., Andrade, B., Panchana, F., Roman, R., Sorroza, L., Trujillo, L.E., Bayot, B., 2022. Microbiome of *Penaeus vannamei* larvae and potential biomarkers associated with high and low survival in shrimp hatchery tanks affected by acute hepatopancreatic necrosis disease. Front. Microbiol. 13, 838640. 10.3389/fmicb.2022.838640.

Ringø, E., 2020. Probiotics in shellfish aquaculture. Aquac. Fish. 5 (1), 1–27. 10.1016/j.aaf.2019.12.001.

Risely, A., 2020. Applying the core microbiome to understand host-microbe systems. J. Anim. Ecol. 89 (7), 1549–1558. 10.1111/1365-2656.13229.

Sanders-Smith, R., Segovia, B.T., Forbes, C., Hessing-Lewis, M., Morien, E., Lemay, M.A., O’Connor, M.I., Parfrey, L.W., 2020. Host-specificity and core taxa of seagrass leaf microbiome identified across tissue age and geographical regions. Front. Ecol. Evol. 8, 605304. 10.3389/fevo.2020.605304.

Shan, X., Li, K., Stadler, P., Borbor, M., Reyes, G., Solorzano, R., Chamorro, E., Bayot, B., Cordero, O.X., 2025. Microbiome determinants of productivity in aquaculture of whiteleg shrimp. Appl. Environ. Microbiol. 91 (5), e02420–24. 10.1128/aem.02420-24.

Shen, W., Le, S., Li, Y., Hu, F., 2016. SeqKit: a cross-platform and ultrafast toolkit for FASTA/Q file manipulation. PLoS One 11 (10), e0163962. 10.1371/journal.pone.0163962.

Sonnenschein, E.C., Jimenez, G., Castex, M., Gram, L., 2021. The *Roseobacter*-group bacterium *Phaeobacter* as a safe probiotic solution for aquaculture. Appl. Environ. Microbiol. 87 (5), e02581–20. 10.1128/AEM.02581-20.

Sonnenschein, E.C., Nielsen, K.F., D’Alvise, P., Porsby, C.H., Melchiorsen, J., Heilmann, J., Kalatzis, P.G., López-Pérez, M., Bunk, B., Spröer, C., 2017. Global occurrence and heterogeneity of the *Roseobacter*-clade species *Ruegeria mobilis*. ISME J. 11 (2), 569–583. 10.1038/ismej.2016.111.

Wang, H., Li, D.H., Wang, J.R., Wang, R., Liang, C.L., Hu, Z., Liu, J.H., 2024. Functional assembly of surface microbiota of *Ulva fasciata* improves nutrient absorption efficiency and growth. Front. Microbiol. 15, 1476073. 10.3389/fmicb.2024.1476073.

Wang, P., Li, S.-P., Yang, X., Si, X., Li, W.-J., Shu, W., Jiang, L., 2023. Spatial scaling of soil microbial co-occurrence networks in a fragmented landscape. mLife 2 (2), 209–215. 10.1002/mlf2.12073.

Wang, Q., Cole, J.R., 2024. Updated RDP taxonomy and RDP Classifier for more accurate taxonomic classification. Microbiol. Resour. Announc. 13 (4), e01063–23. 10.1128/mra.01063-23.

Wang, Y., Wang, K., Huang, L., Dong, P., Wang, S., Chen, H., Lu, Z., Hou, D., Zhang, D., 2020. Fine-scale succession patterns and assembly mechanisms of bacterial community of *Litopenaeus vannamei* larvae across the developmental cycle. Microbiome 8, 106. 10.1186/s40168-020-00879-w.

Wanna, W., Aucharean, C., Jaeram, N., 2025. Analysis of gut microbiota associated with WSSV resistance in *Litopenaeus vannamei*. Mar. Biotechnol. 27, 10. 10.1007/s10126-024-10381-1.

Watts, S.C., Ritchie, S.C., Inouye, M., Holt, K.E., 2019. FastSpar: rapid and scalable correlation estimation for compositional data. Bioinformatics 35 (6), 1064–1066. 10.1093/bioinformatics/bty734.

Wickham, H., Averick, M., Bryan, J., Chang, W., McGowan, L.D.A., François, R., Grolemund, G., Hayes, A., Henry, L., Hester, J., 2019. Welcome to the tidyverse. J. Open Source Softw. 4 (43), 1686. 10.21105/joss.01686.

Wickham, H., François, R., Henry, L., Müller, K., Vaughan, D., 2023. dplyr: A Grammar of Data Manipulation, version 1.1.4 [software]. https://CRAN.R-project.org/package=dplyr.

Yao, Y., Cai, X., Ye, Y., Wang, F., Chen, F., Zheng, C., 2021. The role of microbiota in infant health: from early life to adulthood. Front. Immunol. 12, 708472. 10.3389/fimmu.2021.708472.

Ye, X.S., Chen, M.X., Li, H.Y., He, X.Y., Zhao, Y., 2019. *Halobacteriovorax vibrionivorans* sp. nov., a novel prokaryotic predator isolated from coastal seawater of China. Int. J. Syst. Evol. Microbiol. 69 (12), 3917–3923. 10.1099/ijsem.0.003703.

Yin, X., Ang, L.P., Zhu, R.L., Azarbad, H., Ni, H.H., Chai, M.L., Liu, C., Kong, F., Liu, L.J., Liu, S.L., Ma, Y., Zhou, H.D., Luo, Z.S., He, X.K., Ye, L.X., Li, H., Shu, L., 2025. Host-specific and environmental core bacteria differentially shape the stability and function of the *Sphagnum* phyllosphere. ISME Commun. 5 (1), ycaf221. 10.1093/ismeco/ycaf221.

Yu, J., Jiang, C., Yamano, R., Koike, S., Sakai, Y., Mino, S., Sawabe, T., 2023. Unveiling the early life core microbiome of the sea cucumber *Apostichopus japonicus* and the unexpected abundance of the growth-promoting *Sulfitobacter*. Anim. Microbiome 5, 54. 10.1186/s42523-023-00276-2.

Zhang, F., Chen, H., Yu, J., Chen, R., Zhang, D., Chen, C., Wang, K., 2025. Source tracking of larval bacterial community of Pacific white shrimp across the developmental cycle: ecological insights for microbial management and pathogen prevention in larviculture. Aquaculture 609, 742800. 10.1016/j.aquaculture.2025.742800.

Zhang, S., Sun, C., 2025. Ecological divergence of marine bacteria *Alteromonas mediterranea*. Mol. Phylogenet. Evol. 208, 108359. 10.1016/j.ympev.2025.108359.

Zhang, S., Sun, X., 2022. Core gut microbiota of shrimp function as a regulator to maintain immune homeostasis in response to WSSV infection. Microbiol. Spectr. 10 (2), e02465–21. 10.1128/spectrum.02465-21.

Zhang, Z., Yang, Q., Liu, H., Jin, J., Yang, Y., Zhu, X., Han, D., Zhou, Z., Xie, S., 2025. Potential functions of the gut microbiome and modulation strategies for improving aquatic animal growth. Rev. Aquac. 17 (1), e12959. 10.1111/raq.12959.

Zhou, K.-M., Liu, P.-P., Yao, J.-Y., Vasta, G.R., Wang, J.-X., Wang, X.-W., 2025. Shrimp intestinal microbiota homeostasis: dynamic interplay between the microbiota and host immunity. Rev. Aquac. 17 (1), e12986. 10.1111/raq.12986.

Zhou, X., Shen, X., Johnson, J.S., Spakowicz, D.J., Agnello, M., Zhou, W., Avina, M., Honkala, A., Chleilat, F., Chen, S.J., 2024. Longitudinal profiling of the microbiome at four body sites reveals core stability and individualized dynamics during health and disease. Cell Host Microbe 32 (4), 506–526. e9. 10.1016/j.chom.2024.02.012.

