## Supplementary information for "Early-life core microbiota are associated with postlarval survival of *Penaeus vannamei*: A meta-analysis"


The supplementary information contains:

- Supplementary Methods
- Supplementary Results
- Supplementary Figures S1-S8
- Supplementary Tables S1-S4

### Supplementary Methods

#### SM1. Additional larviculture cycles conducted in this study

Two additional larviculture cycles, as referred to as Studies 12 and 13 (Table S1), were respectively conducted at the Yongxing Base (27.859° N, 120.845° E) and the Qingjiang Base (28.280° N, 121.110° E) of Zhejiang Mariculture Research Institute, Wenzhou, Zhejiang, China. The larviculture cycle of Study 12 lasted for 21 days post hatching (dph) (a prolonged cycle due to reduced water temperature resulting from a heating system malfunction at *zoea* III stage and *postlarvae* I stage) in June 2023, while that of Study 13 lasted for 16 dph in June 2024. Both cycles covered the developmental stages from *nauplius* I to *postlarvae* V. The cycle of Study 12 involved six uniform ponds (7 m × 5 m × 2 m) culturing larvae at an initial density of $1.96 \times 10^5$ nauplii/$m^3$. The six ponds were randomly assigned to either the control group or the probiotic-supplemented group, with three replicates per for each group. Live microalgae (*Arthrospira* sp.) were provided as the initial feed from the *nauplius* VI to the *mysis* I stage. From the *zoea* I stage onward, larvae were continuously fed with shrimp flakes and *Artemia* (frozen ones prior to *mysis* and live ones thereafter). In the probiotic-supplemented group, a *Ruegeria profundi* strain was co-supplied with dry feeds at a concentration of $10^7$ CFU/g twice daily. The cycle of Study 13 involved eight ponds (4 m × 5 m × 1.6 m) culturing larvae of two shrimp strains at an initial density of $2.8 \times 10^5$ nauplii/$m^3$, with four ponds allocated to each shrimp strain. Live microalgae (*Chaetoceros* sp.) were provided as the initial feed from the *nauplius* VI stage to the *zoea* III stage. From the *zoea* I stage onward, larvae were continuously fed with shrimp flakes, *Artemia* (frozen ones supplied from *zoea* II to *mysis* III, while live ones supplied from the *postlarvae* I stage onward). Final larval (postlarval) survival rates were recorded at the end of both larviculture cycles (*postlarvae* V).

Laval and/or rearing water samples were collected from each pond across host developmental stages (Study 12: *nauplius* I, *zoea* I, *zoea* II, *zoea* III, *mysis* I, *mysis* III, *postlarvae* I, *postlarvae* III, and *postlarvae* V; Study 13: *nauplius* I, *zoea* I, *zoea* III, *mysis* I, *mysis* III, *postlarvae* I, and *postlarvae* V) as previously described (Zhang et al., 2025). In total, 54 larval samples and 54 water filters from Study 12, as well as 56 larval samples from Study 13, were used for 16S rRNA gene amplicon sequencing.

**SM2. DNA extraction, 16S rRNA gene amplification, and Illumina sequencing**

Total DNA in the larval samples and water filters was extracted using a QIAamp® DNA Stool Mini Kit (QIAGEN, Germany) and a SPINeasy DNA Kit for Soil (MPBIO, USA), respectively. The V4 region of 16S rRNA genes was amplified using the primers set 515FY (5'-GTGYCAGCMGCCGCGGTAA-3') and 806RB (5'-GGACTACNVGGGTWTCTAAT-3') with dual barcodes (Apprill et al., 2015; Parada et al., 2016). An amount of 50 ng purified DNA template from each sample was amplified with a 50-μl reaction system. The PCR procedure of Study 12 consisted of an initial denaturation at 95 °C for 2 min, followed by 30 cycles of denaturation at 95 °C for 15 s, annealing at 53 °C for 15 s, and extension at 72 °C for 15 s, with a final extension at 72 °C for 5 min, while that of Study 13 involved an initial denaturation at 94 °C for 5 min, followed by 31 cycles of denaturation at 94 °C for 30 s, annealing at 53 °C for 30 s, and extension at 72 °C for 30 s, with a final extension at 72 °C for 8 min. The PCR products were purified with an E.Z.N.A.® Gel Extraction Kit (Omega, USA). The library was prepared with an ALFA-SEQ DNA Library Prep Kit (FINDROP, China), quantified using a Qubit 4.0 fluorometer (Invitrogen, USA), and then sequenced on an Illumina NovaSeq 6000 platform (Illumina, USA; Study 12) or an Illumina NovaSeq X Plus platform (Illumina, USA; Study 13) at MAGIGENE Co., Ltd., Guangzhou, China.

**SM3. General statistical analysis**

All statistical analyses and data visualization were performed in the R environment (v4.4.2) unless otherwise indicated. The geographic distribution of samples was visualized using the R package 'maps' (Becker et al., 2025). Sankey diagram was generated to show the flow of samples through different attributes using the R package 'networkD3' (Allaire et al., 2025). Comparisons of larval and water prokaryotic alpha diversity across geographic locations, or among samples from the same location but different larviculture batches at a given developmental stage were performed with Mann-Whitney *U* test using the R package 'rstatix' (Kassambara, 2025). Analysis of Variance (ANOVA) was applied to assess the influence of sample type, developmental stage, geographic location, and their interactions on alpha-diversity indices using the R package 'car' (Fox and Weisberg, 2019). For the 265 paired larva-water samples, linear models were used to examine the relationships between prokaryotic alpha diversity in the rearing water and that in the larvae. Principal Coordinates Analysis (PCoA) was performed to visualize the compositional variation of prokaryotic communities across larval and water samples, geographic locations, and larval developmental stages using the R packages 'vegan' and 'phyloseq' (McMurdie and Holmes, 2013). Permutational Multivariate Analysis of Variance (PERMANOVA) and Analysis of Similarity (ANOSIM) were performed to assess the effects of sample type, developmental stage, geographic location, and their interactions on prokaryotic community composition using the package 'vegan'. The relationship between community similarity and geographic distance between locations was analyzed using the R package 'geosphere' (Hijmans et al., 2026).

**Supplementary Results**

**SR1. Taxonomic composition of prokaryotic communities in rearing water**

As observed in larval samples, prokaryotic communities in rearing water were overwhelmingly dominated by bacteria, accounting for 99.90% of the entire water dataset. At higher taxonomic ranks, water microbial communities were primarily composed of *Alphaproteobacteria*, *Bacteroidota*, *Actinomycetota*, and *Gammaproteobacteria*, although their relative abundances varied across larval developmental stages and among geographic locations (only samples from China were available) (Fig. S4). *Alphaproteobacteria* predominated across all sites, whereas the contributions of *Bacteroidota*, *Gammaproteobacteria*, and *Actinomycetota* differed markedly among locations. In addition, *Cyanobacteriota* occasionally became dominant in Wenzhou, particularly during the *nauplius* and *zoea* stages, indicating pronounced stage-specific shifts in water microbial composition. At the family level, rearing water prokaryotic communities were consistently dominated by a small number of bacterial families, with *Roseobacteraceae* (mean relative abundance: 36.07%), *Flavobacteriaceae* (14.89%), *Lewinellaceae* (8.34%), *Cyclobacteriaceae* (5.27%), and *Microbacteriaceae* (2.95%) together accounting for 67.52% of the entire water microbiota dataset. Despite this overall consistency, pronounced location- and stage-dependent patterns were evident in family-level composition. Specifically, rearing water in Wenzhou were characterized by relatively higher abundances of *Roseobacteraceae*, *Lewinellaceae*, and *Flavobacteriaceae*, whereas those in Shenzhen were strongly dominated by *Roseobacteraceae* and *Flavobacteriaceae*, accompanied by elevated *Microbacteriaceae*. In Zhanjiang, *Roseobacteraceae* and F*lavobacteriaceae* likewise dominated, with moderate contributions from *Lewinellaceae* and *Balneolaceae*. In contrast, rearing water in Wenchang exhibited a distinct composition, marked by a high relative abundance of *Cyclobacteriaceae* in addition to *Roseobacteraceae* and *Flavobacteriaceae*. Collectively, these results indicate that, although a conserved set of dominant bacterial families prevailed in rearing water, their relative abundances varied substantially across geographic locations and larval developmental stages.

**Supplementary Figures**

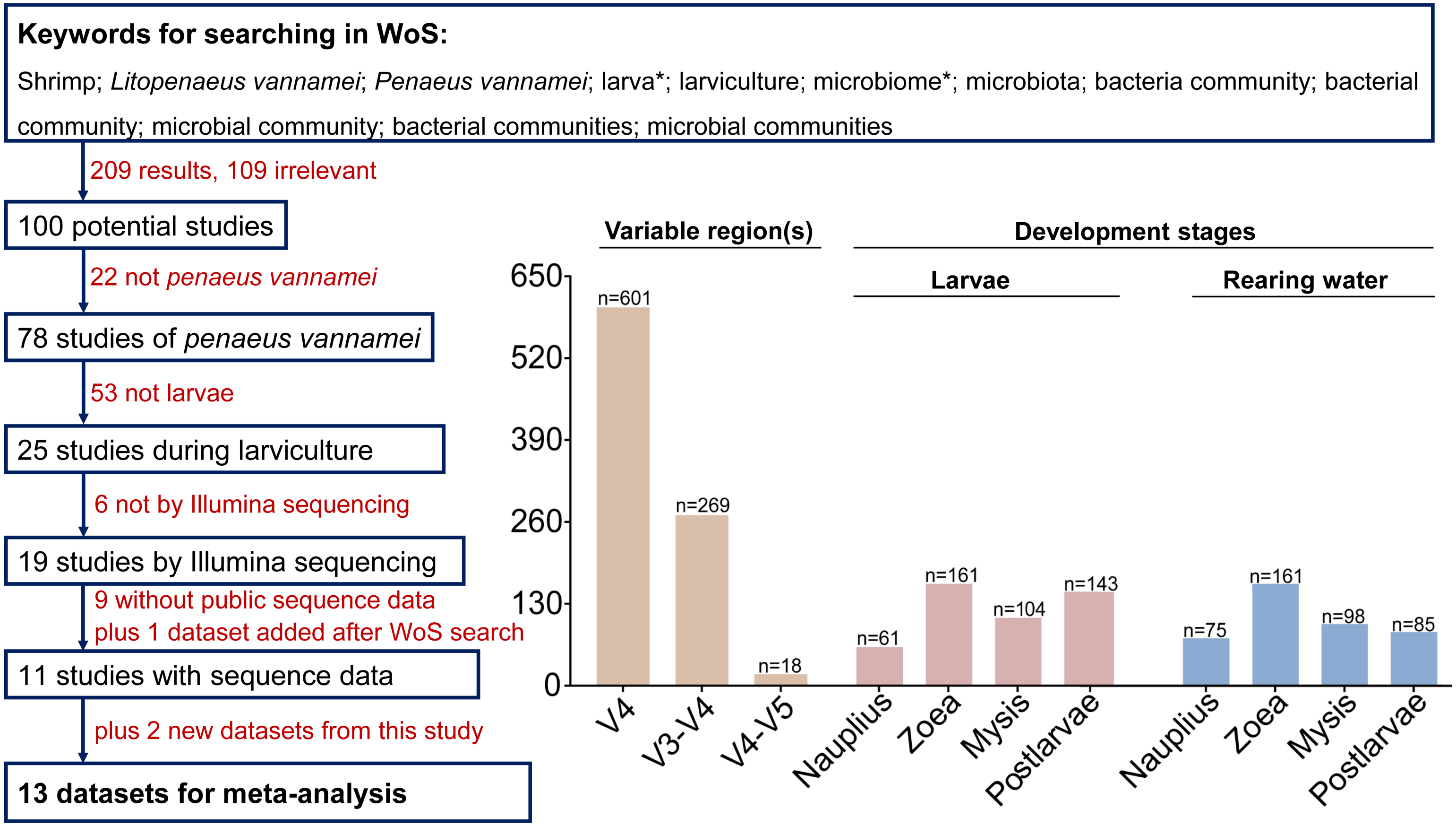


**Figure S1.** Literature screening workflow and sample distribution of 16S rRNA gene microbiota regarding *Penaeus vannamei* larviculture. The flowchart (left) showing the screening procedure from 209 initial Web of Science (WoS) search (published before 12 November 2024). A total of 13 datasets covering variable regions that include the V4 region were selected. These comprised 11 datasets from peer-reviewed publications (including one published by our laboratory after the WoS search, referred to as Study 11) and two previously unpublished datasets generated by our laboratory, referred to as Study 12 and Study 13 (Table S1). The bar plots (right) showing the number of samples for different variable regions of the 16S rRNA gene (including V4, V3-V4, and V4-V5), as well as of both larvae and rearing water across developmental stages, including *nauplius*, *zoea*, *mysis*, and *postlarvae*. Samples with mixed or ambiguous information of developmental stages were excluded from the final dataset for analysis.

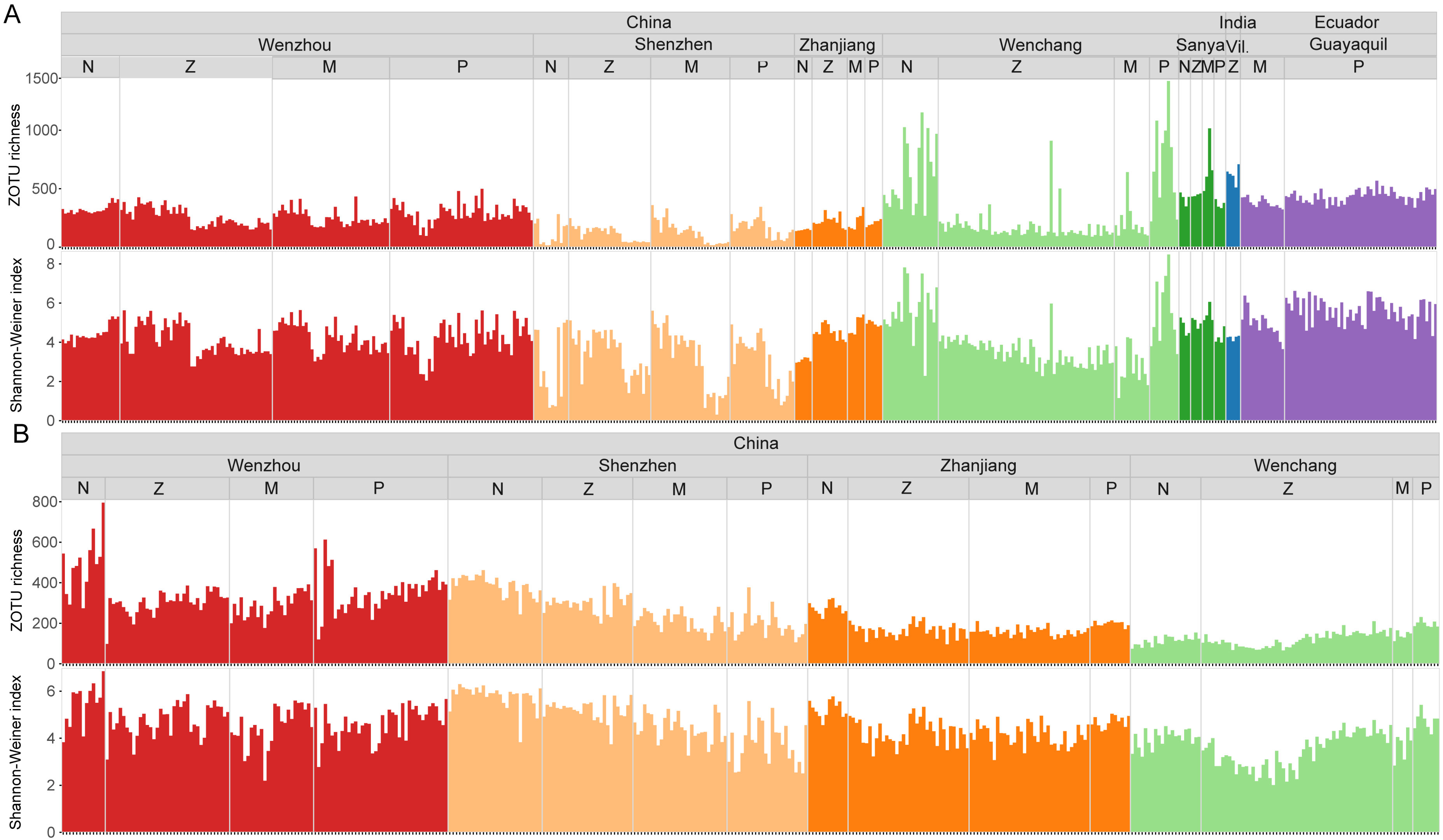


**Figure S2.** Alpha diversity of prokaryotic communities in shrimp larvae and rearing water. Variation in prokaryotic alpha-diversity indices in larvae (A) and water (B) across geographic locations and larval developmental stages. N, *nauplius*; Z, *zoea*; M, *mysis*; P, *postlarvae*. Vil., Viluppuram.

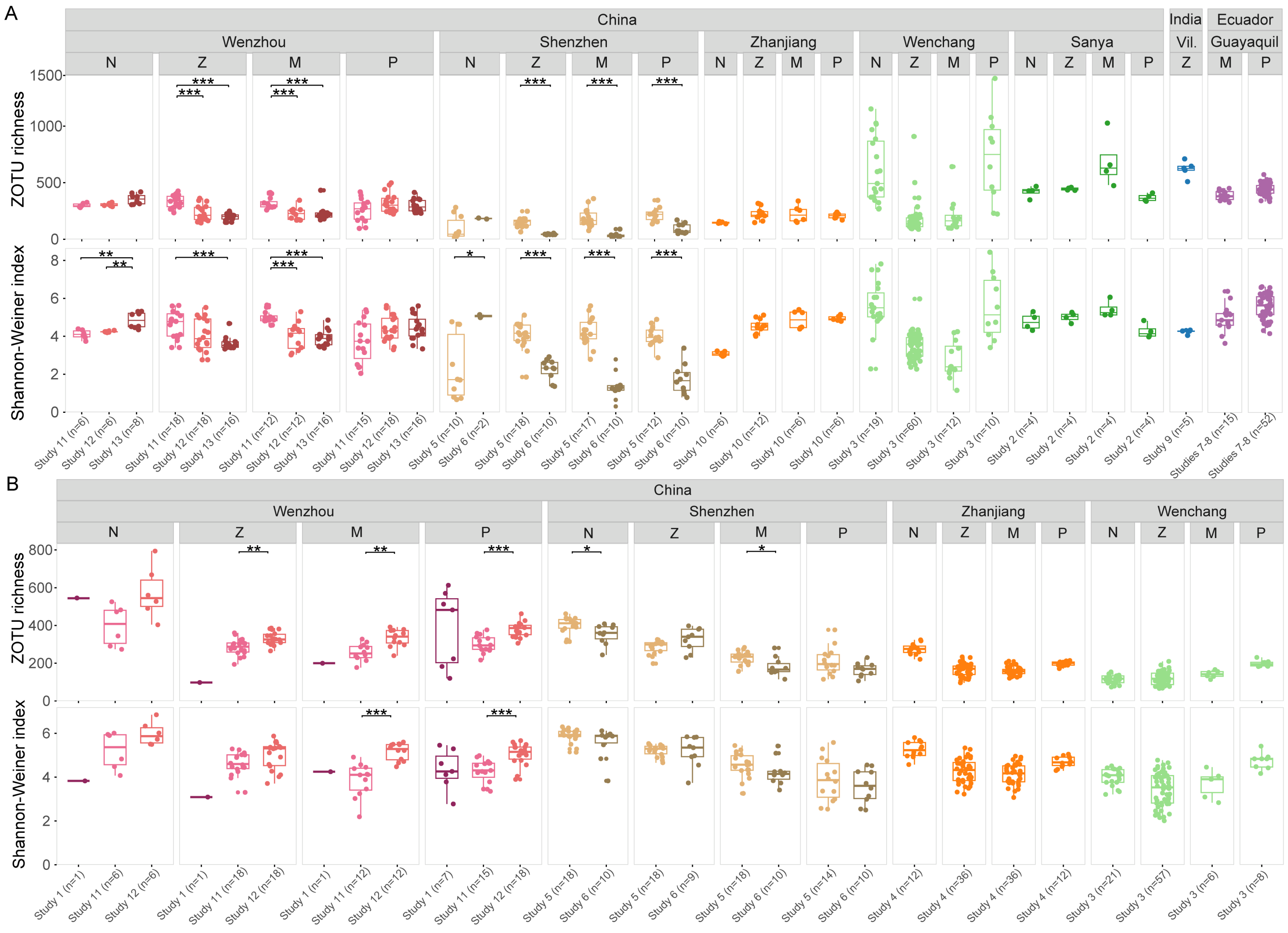


**Figure S3.** Comparison of alpha-diversity indices of prokaryotic communities in larvae and rearing water. Differences in prokaryotic alpha-diversity indices in larvae (A) and rearing water (B) among different larviculture cycles (batches) within the same location or larval developmental stage. The boxes illustrate the median and interquartile range, and the whiskers extend from the minimum to the maximum values. Asterisks indicate significant differences between larviculture batches/studies (if any) at the same location and developmental stage, as determined by the Mann-Whitney *U* test with false discovery rate (FDR) correction ($^{*}P < 0.05$, $^{**}P < 0.01$, $^{***}P < 0.001$). N, *nauplius*; Z, *zoea*; M, *mysis*; P, *postlarvae*. Vil., Viluppuram.

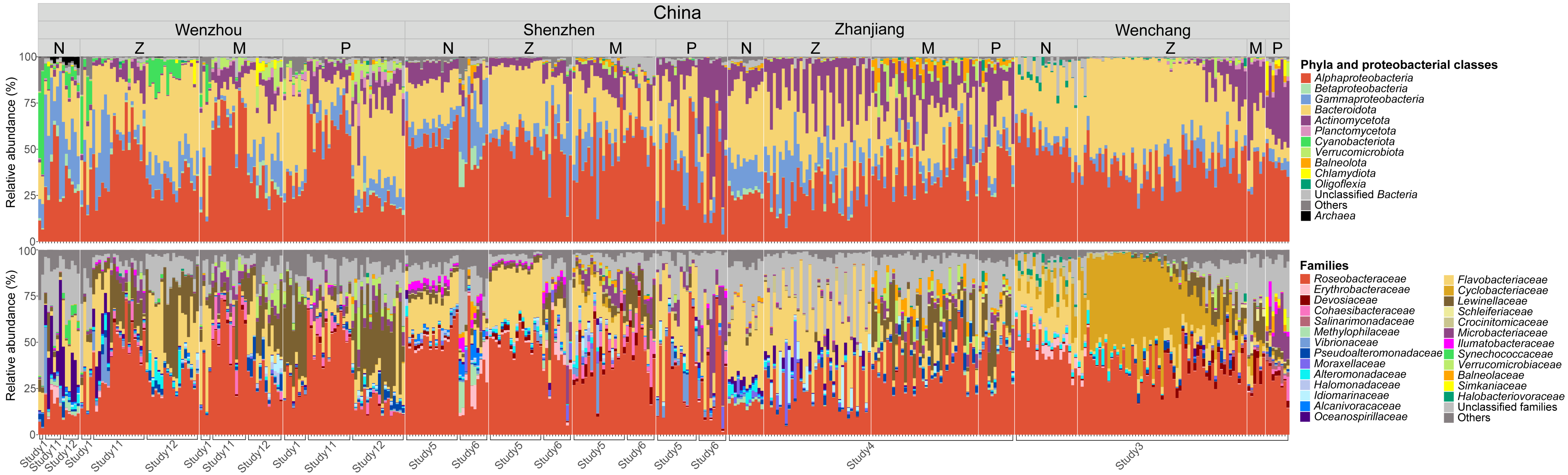


**Figure S4.** Taxonomic distribution of rearing water prokaryotic communities across geographic locations and larval developmental stages. The upper and bottom panels show phylum- and family-level taxonomic composition, respectively, with *Proteobacteria* (now *Pseudomonadota*) further classified at the class level. Taxa with average relative abundance ≥ 2% at least at one developmental stage in one city are shown. N, *nauplius*; Z, *zoea*; M, *mysis*; P, *postlarvae*.

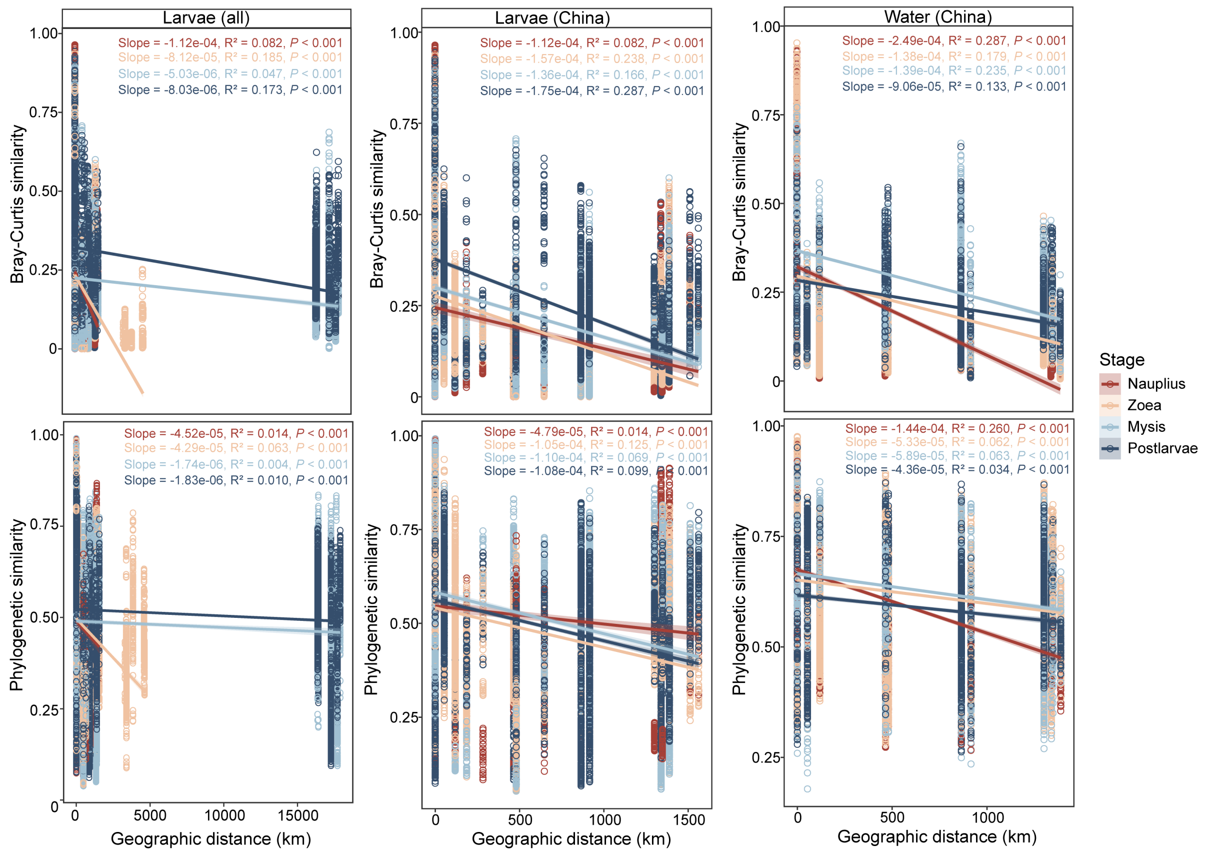


**Figure S5.** Relationships of taxonomic and phylogenetic similarity of larval and rearing-water prokaryotic communities with geographic distance. Taxonomic and phylogenetic similarities were estimated as Bray–Curtis similarity and 1 − weighted UniFrac distance, respectively. Linear regressions were performed for each developmental stage using all larval samples (left panels), larval samples from China (middle panels), and rearing water samples (China only; right panels).

A

| Occupancy (%) | Number of common core ZOTUs at each stage | | | | Number of pan-common core ZOTUS |
|---|---|---|---|---|---|
| | Nauplius | Zoea | Mysis | Postlarvae | |
| **50** | 111 | 22 | 80 | 104 | **163** |
| **60** | 65 | 18 | 52 | 64 | **95** |
| **70** | 41 | 9 | 34 | 40 | **58** |
| **80** | 24 | 7 | 22 | 26 | **35** |
| **90** | 7 | 0 | 10 | 17 | **20** |
| **100** | 0 | 0 | 1 | 4 | **5** |

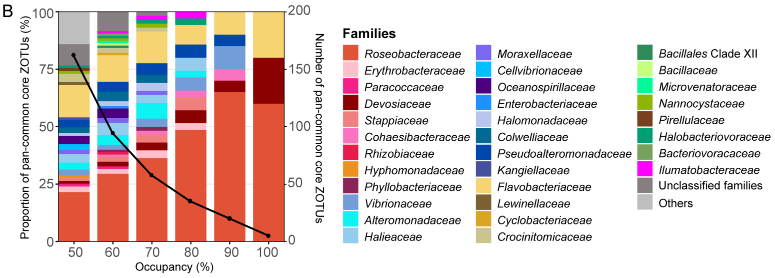


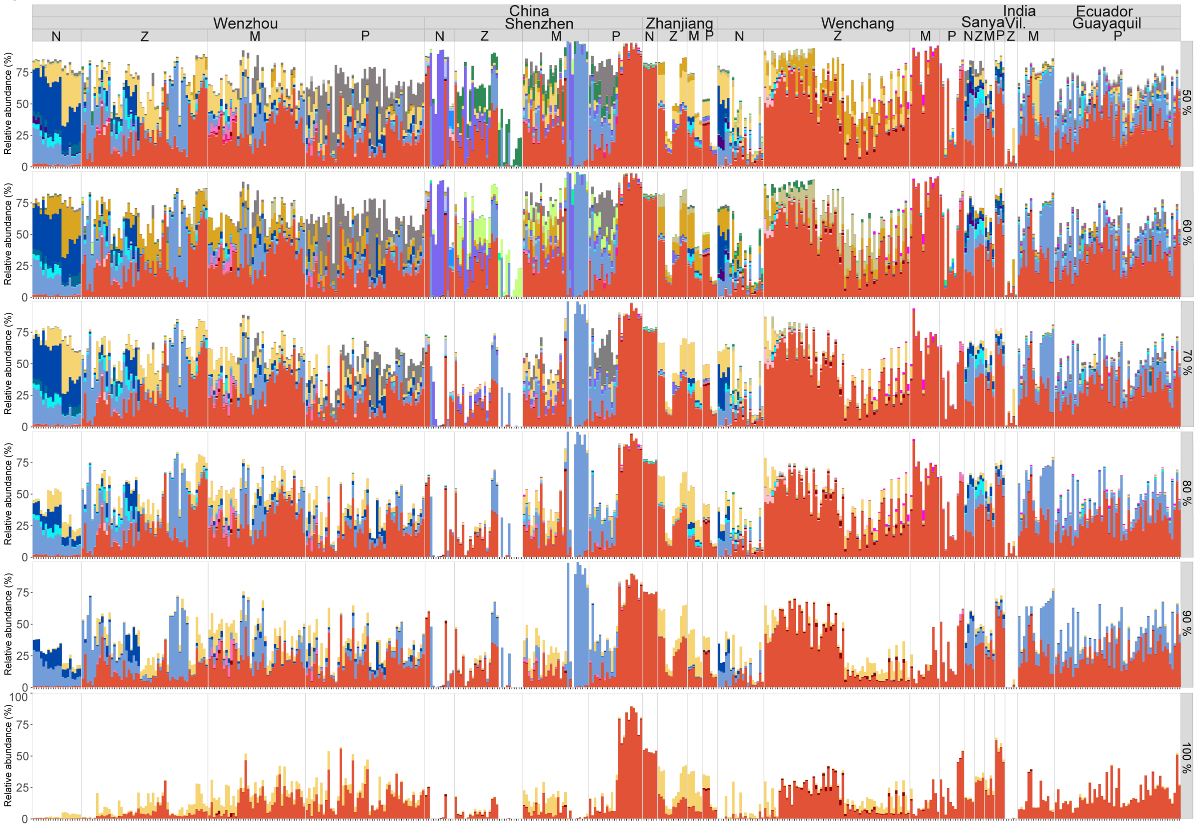


**Figure S6.** Robustness test of occupancy thresholds for identifying common core ZOTUs in shrimp larvae. Number of common core ZOTUs at each developmental stage under varying occupancy thresholds ranging from 50% to 100% at 10% intervals (A). Proportions of pan-common core ZOTUs classified at the family level across different thresholds (the families with proportions ≥ 1% are shown) (B). Family-level taxonomic composition of larval pan-common core ZOTUs (identified under different thresholds) across geographic locations and developmental stages (C). N, *nauplius*; Z, *zoea*; M, *mysis*; P, *postlarvae*. Vil., Viluppuram.

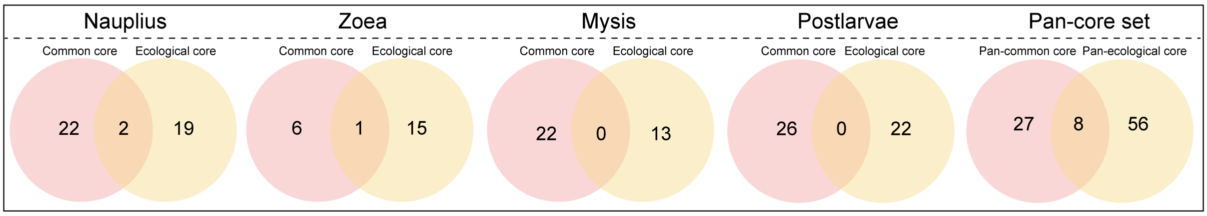


**Figure S7.** The overlap of common core ZOTUs and ecological core ZOTUs in the larval prokaryotic community. Venn diagrams illustrates their overlaps at each developmental stage and during the entire developmental cycle (as the pan-core sets).

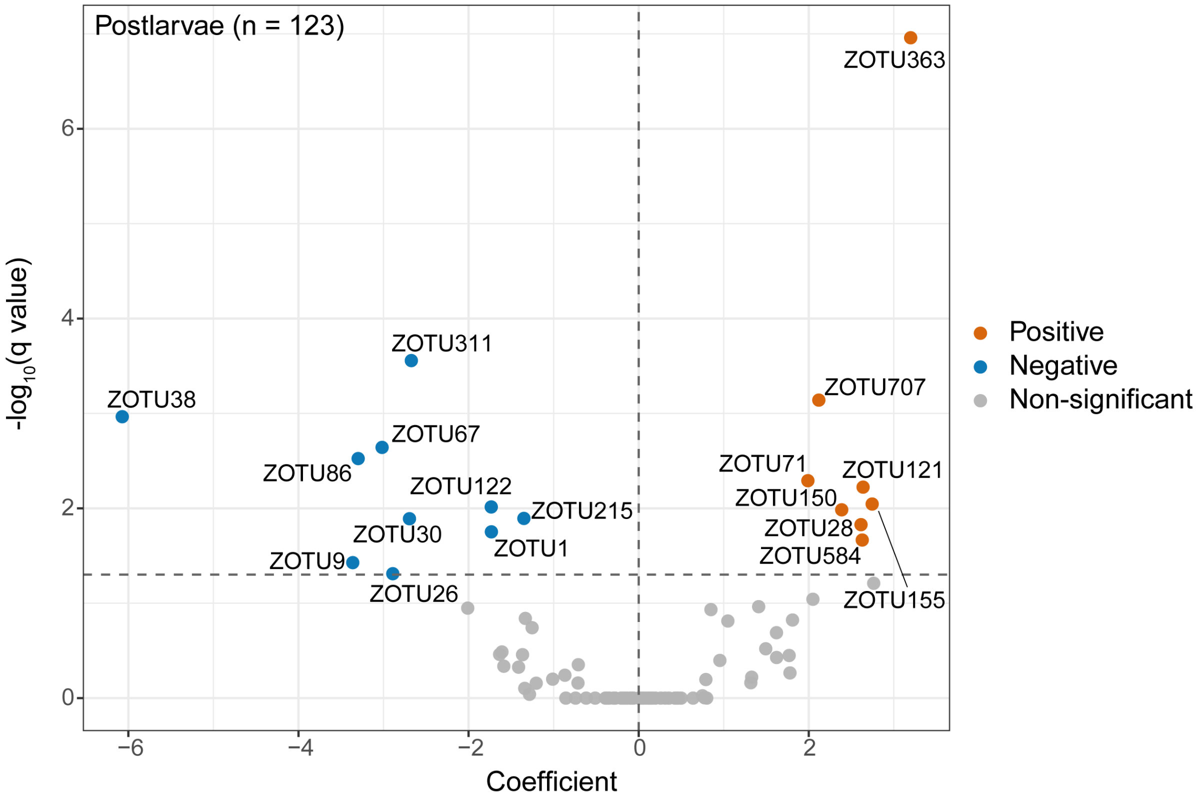


**Figure S8.** Volcano plot showing multivariable linear correlations between pan core ZOTUs and postlarval survival rate. The pan core ZOTUs include pan-common core, temporal core, and pan-ecological core ZOTUs as identified in Figs. 5 and 6. Of 91 pan core ZOTUs in total, ZOTU396 was excluded due to its absence in the larval samples with postlarval survival rate data. Orange and blue dots denote ZOTUs showing significantly positive and negative associations with postlarval survival rate, respectively ($q < 0.05$ after Benjamini-Hochberg correction), whereas grey dots denote no significance.

**Supplementary Tables**

**Table S1.** Key information of the datasets used for the meta-analysis.

| Study ID | Location | Sampling time | Checked sample number | Developmental stages | Experimental group setting or sampling scheme | Health status | Initial larval density (nauplii/m³) | Mouth-opening feeds | Reference |
|---|---|---|---|---|---|---|---|---|---|
| 1* | Wenzhou, Zhejiang, China (28.016°N, 120.642°E) | Undescribed | Water: 18 | N; Z; M; P | Left pool; Right pool | Undescribed | $2.5\times10^5$ | Unicellular algae; Plant debris | (Yan et al., 2020) |
| 2* | Sanya, Hainan, China (18.617°N, 109.950°E) | Nov.-Dec. 2017 | Larvae: 16 | $N_V$; $Z_{II}$; $M_{II}$; $P_{II}$ | 4 ponds: D801-D804 | Free from specific pathogens (EHP, $V_{AHPND}$, DIV1, WSSV, IHHNV) | $1.17\times10^6$ | Live algae: *Thalassiosira pseudonana* | (Wang, H. et al., 2020) |
| 3 | Wenchang, Hainan, China (20.148°N, 110.687°E) | Aug. 2016 | Larvae: 101; Water: 92 | N; $Z_{(I, II, III)}$; M; P | Three random ponds for all-time sampling (26 hourly time points); Other three ponds were sampled at one time point of each stage (except two time points at stages $Z_{III}$, M, and P) | Undescribed | $2.69\times10^5$ | Live microalgae: *Chaetoceros* sp., *Thalassiosira* sp.; Shrimp flakes | (Wang, Y. et al., 2020) |
| 4* | Zhanjiang, Guangdong, China (21.154°N, 110.298°E) | Undescribed | Water: 96 | $N_V$; $Z_{(I, II, III)}$; $M_{(I, II, III)}$; $P_{II}$ | 12 ponds: CX1-CX12 | CX6: A mass mortality at stage $M_{II}$; Others: No visible symptoms of disease | $2\times10^5$ | Commercial spirulina powder; Shrimp flakes | (Xue et al., 2020) |

| Study ID | Location | Sampling time | Checked sample number | Developmental stages | Experimental group setting or sampling scheme | Health status | Initial larval density (nauplii/m³) | Mouth opening feeds | Reference |
|---|---|---|---|---|---|---|---|---|---|
| 5* | Shenzhen, Guangdong, China (22.557°N, 114.539°E) | Undescribed | Larvae: 57; Water: 68 | $N_{VI}$; $Z_I$; $M_I$; $P_I$ | Control; +Effective microorganisms; +*Bacillus* sp.; +*Lactobacillus* sp.; +Photosynthetic bacteria; +*Saccharomyces* sp. | Undescribed | $3\times10^5$ | Undescribed | (Wang, R. et al., 2020) |
| 6* | Shenzhen, Guangdong, China (22.557°N, 114.539°E) | Undescribed | Larvae: 32; Water: 40 | $N_{VI}$; $Z_I$; $M_I$; $P_I$ | Control; +$ClO_2$; +HCHO; +CaClO; +$I_2$ | Undescribed | $2\times10^5$ | Spirulina powder; Prawn crackers | (Duan et al., 2020) |
| 7* | Guayaquil, Guayas, Ecuador (2.148°S, 79.965°W) | Jan.-Mar. 2021 | Larvae: 47 | $M_{III}$; $P_{(IV, VII, X)}$ | High survival group (> 60 %); Low survival group (< 60 %) | AHPND | NA | Mixed commercial feeds | (Reyes et al., 2022) |
| 8* | Guayaquil, Guayas, Ecuador (2.148°S, 79.965°W) | Jan.-Mar. 2021 | Larvae: 20 | $M_{III}$; $P_{(IV, VII, X)}$ | AHPND group (low survival: < 60 %); Zoea-II syndrome group | AHPND; Zoea-II syndrome | NA | Mixed commercial feeds | (Reyes et al., 2023) |
| 9 | Viluppuram, Tamil Nadu, India (12.19°N, 79.92°E) | Undescribed | Larvae: 5 | $Z_I$ | Control; (2, 4, 6, 8) days post-infection | Zoea-II syndrome | NA | Undescribed | (Thangaraj et al., 2023) |

| Study ID | Location | Sampling time | Checked sample number | Developmental stages | Experimental group setting or sampling scheme | Health status | Initial larval density (nauplii/m³) | Mouth opening feeds | Reference |
|---|---|---|---|---|---|---|---|---|---|
| 10* | Zhanjiang, Guangdong, China (21.154°N, 110.298°E) | Undescribed | Larvae: 30 | $N_V$; $Z_{II}$; $M_I$; $P_I$ | DNA extracted by different kits: Bacterial DNA Kit; Mollusc DNA Kit; Stool DNA Kit; Tissue DNA Kit | No visible symptoms of disease | NA | Undescribed | (Xue et al., 2018) |
| 11 | Wenzhou, Zhejiang, China (27.859°N, 120.845°E) | Jun. 2022 | Larvae: 51; Water: 51 | $N_I$; $Z_{(I, II, III)}$; $M_{(I, 9dph)}$; $P_{(I, 13dph, V)}$ | Two shrimp strains: Stress-resistant; Fast-growing | Near extinction in one of the fast-growing ponds by day 10; Others: No visible symptoms of disease | $1.43\times10^5$ | Live microalgae: *Chaetoceros* sp., *Isochrysis* sp., *Dicrateria* sp. | (Zhang et al., 2025) |
| 12# | Wenzhou, Zhejiang, China (27.859°N, 120.845°E) | Jun. 2023 | Larvae: 54; Water: 54 | $N_I$; $Z_{(I, II, III)}$; $M_{(I, III)}$; $P_{(I, III, V)}$ | Control; +*Ruegeria profundi* strain | Developmental delay at stages $Z_{III}$ and $P_I$ | $1.96\times10^5$ | Live algae: *Arthrospira* sp.; Shrimp flakes | This study (a) |
| 13# | Wenzhou, Zhejiang, China (28.282°N, 121.109°E) | Jun. 2024 | Larvae: 56 | $N_I$; $Z_{(I, III)}$; $M_{(I, III)}$; $P_{(I, V)}$ | Two shrimp strains: Stress-resistant; Fast-growing | No visible symptoms of disease | $2.8\times10^5$ | Live algae: *Chaetoceros* sp. | This study (b) |

Well symbol (#) indicates two previously unpublished datasets generated by our laboratory. Asterisks (*) indicates those studies without disclosed geographic coordinates in the text, for which the geographic coordinates of the institution hosting the experimental site were used instead. N, *nauplius*; Z, *zoea*; M, *mysis*; P, *postlarvae*. Subscript letters following the abbreviations of developmental stages represent their respective sub-stages. AHPND, acute hepatopancreatic necrosis disease; EHP, *Enterocytozoon hepatopenaei*; $V_{AHPND}$, Acute hepatopancreatic necrosis disease-causing *Vibrio*; DIV1, Decapod iridescent virus 1; WSSV, White spot syndrome virus; IHHNV, Infectious hypodermal and hematopoietic necrosis virus; NA, not available.

**Table S2.** DNA extraction protocols, 16S rRNA gene primers, and sequence-trimming approaches used to generate the datasets.

| Study ID | DNA extraction protocol | 16S region | Primer set | Accession number | Sequence-trimming method | Average pre-trim sequence length (nt) | Average post-trim sequence length (nt) | Average sequence recovery rate | Average post-trim read depth | Reference |
|---|---|---|---|---|---|---|---|---|---|---|
| 1 | **Water:** Water DNA Kit (OMEGA Biotek, USA) | V4-V5 | 515F (5'-GTGCCAGCMGCCGCGGTAA-3'); 909R (5'-CCCCGYCAATTCMTTTRAGT-3') | PRJNA560666 | e-PCR (USEARCH script *search_pcr2*) | 418.4 | 253 | 95.91% | 28,574 | (Yan et al., 2020) |
| 2 | **Larvae:** Standard CTAB method | V4 | 515F (5'-GTGCCAGCMGCCGCGGTAA-3'); 806R (5'-GGACTACHVGGGTWTCTAAT-3') | PRJNA547842 | Already processed | 250.7 | 250.7 | 100% | 87,315 | (Wang, H. et al., 2020) |
| 3 | **Larvae:** QIAamp® DNA Stool Mini Kit (QIAGEN, Germany); **Water:** Power Soil® DNA Kit (MOBIO, USA) | V4 | 515F-Y (5'-GTGYCAGCMGCCGCGGTAA-3'); 806R-B (5'-GGACTACNVGGGTWTCTAAT-3') | PRJDB9538 | Trimming the residual reverse primer (USEARCH script *fastx_truncate*) | 273 | 253 | 100% | 36,938 | (Wang, Y. et al., 2020) |
| 4 | **Water:** Water DNA Kit (OMEGA, USA) | V4 | 515F (5'-GTGCCAGCMGCCGCGGTAA-3'); 806R (5'-GGACTACHVGGGTWTCTAAT-3') | CRA001512 | Already processed | 253 | 253 | 100% | 34,020 | (Xue et al., 2018) |

| Study ID | DNA extraction protocol | 16S region | Primer set | Accession number | Sequence-trimming method | Average pre-trim sequence length (nt) | Average post-trim sequence length (nt) | Average sequence recovery rate | Average post-trim read depth | Reference |
|---|---|---|---|---|---|---|---|---|---|---|
| 5 | **Larvae &Water:** Mabio DNA Mini Kit (Jirui Gene Technology, China) | V3-V4 | 341F (5'-CCTACGGRRBGCASCAGKVRVGAAT-3'); 806R (5'-GGACTACNVGGGTWTCTAATCC-3') | PRJNA614602 | e-PCR (USEARCH script *search_pcr2*) | 446.1 | 253 | 56.97% | 57,326 | (Wang, R. et al., 2020) |
| 6 | **Larvae &Water:** DNA Isolation Kit (QIAGEN, USA) | V3-V4 | 341F (5'-CCTACGGRRBGCASCAGKVRVGAAT-3'); 806R (5'-GGACTACNVGGGTWTCTAATCC-3') | PRJNA600559 | e-PCR (USEARCH script *search_pcr2*) | 447.7 | 253 | 52.06% | 23,046 | (Duan et al., 2020) |
| 7 | **Larvae:** ZymoBIOMICS DNA Microprep Isolation Kit (Zymo Research, USA) | V3-V4 | 341F (5'-CCTACGGGAGGCAGCAG-3'); 806R (5'-GGACTACHVGGGTWTCTAAT-3') | PRJNA800805 | Trimming according to the V4 forward primer with Cutadapt | 414.9 | 254 | 100% | 121,490 | (Reyes et al., 2022) |
| 8 | **Larvae:** ZymoBIOMICS DNA Microprep Isolation Kit (Zymo Research, USA) | V3-V4 | 341F (5'-CCTACGGGAGGCAGCAG-3'); 806R (5'-GGACTACHVGGGTWTCTAAT-3') | PRJNA870476 | Trimming according to the V4 forward primer with Cutadapt | 416.6 | 253.9 | 100% | 122,288 | (Reyes et al., 2023) |

| Study ID | DNA extraction protocol | 16S region | Primer set | Accession number | Sequence-trimming method | Average pre-trim sequence length (nt) | Average post-trim sequence length (nt) | Average sequence recovery rate | Average post-trim read depth | Reference |
|---|---|---|---|---|---|---|---|---|---|---|
| 9 | **Larvae:** CTAB and Phenol-chloroform method followed by RNase A treatment | V3-V4 | 341F (5'-GCCTACGGGNGGCWGCAG-3'); 806R (5'-ACTACHVGGGTATCTAATCC-3') | PRJNA832951 | e-PCR (USEARCH script *search_pcr2*) | 459.8 | 253 | 95.91% | 328,488 | (Thangaraj et al., 2023) |
| 10 | **Larvae:** Bacterial DNA Kit (OMEGA, USA); Mollusc DNA Kit (OMEGA, USA); Stool DNA Kit (OMEGA, USA); Tissue DNA Kit (OMEGA, USA) | V4 | 515F (5'-GTGCCAGCMGCCGCGGTAA-3'); 806R (5'-GGACTACHVGGGTWTCTAAT-3') | CRA000198 | Already processed | 253 | 253 | 100% | 45,605 | (Xue et al., 2018) |
| 11 | **Larvae:** QIAamp® Fast DNA Stool Mini Kit (QIAGEN, Germany); **Water:** SPINeasy DNA Kit for Soil (MPBIO, USA) | V4 | 515F-Y (5'-GTGYCAGCMGCCGCGGTAA-3'); 806R-B (5'-GGACTACNVGGGTWTCTAAT-3') | PRJNA1188736 | e-PCR (USEARCH script *search_pcr2*) | 292 | 253 | 100% | 145,933 | (Zhang et al., 2025) |

| Study ID | DNA extraction protocol | 16S region | Primer set | Accession number | Sequence-trimming method | Average pre-trim sequence length (nt) | Average post-trim sequence length (nt) | Average sequence recovery rate | Average post-trim read depth | Reference |
|---|---|---|---|---|---|---|---|---|---|---|
| 12# | **Larvae:** QIAamp® Fast DNA Stool Mini Kit (QIAGEN, Germany); **Water:** SPINeasy DNA Kit for Soil (MPBIO, USA) | V4 | 515F-Y (5'-GTGYCAGCMGCCGCGGTAA-3'); 806R-B (5'-GGACTACNVGGGTWTCTAAT-3') | PRJNA1381294 | e-PCR (USEARCH script *search_pcr2*) | 292 | 253 | 99.99% | 118,497 | This study (a) |
| 13# | **Larvae:** QIAamp® Fast DNA Stool Mini Kit (QIAGEN, Germany) | V4 | 515F-Y (5'-GTGYCAGCMGCCGCGGTAA-3'); 806R-B (5'-GGACTACNVGGGTWTCTAAT-3') | PRJNA1381294 | e-PCR (USEARCH script *search_pcr2*) | 291.7 | 252.7 | 99.97% | 149,291 | This study (b) |

Well symbol (#) indicates two previously unpublished datasets generated by our laboratory.

**Table S3.** Two-way and three-way repeated measures of Analysis of Variance (ANOVA) on prokaryotic alpha diversity.

| Sample type | Factor | ZOTU richness | | Shannon-Wiener index | |
|---|---|---|---|---|---|
| | | F | *P* | F | *P* |
| Larva | Location | 56.76 | < **0.001** | 33.29 | < **0.001** |
| | Stage | 39.61 | < **0.001** | 7.185 | < **0.001** |
| | Location × Stage | 20.91 | < **0.001** | 10.28 | < **0.001** |
| Water | Location | 287.1 | < **0.001** | 68.19 | < **0.001** |
| | Stage | 70.10 | < **0.001** | 41.35 | < **0.001** |
| | Location × Stage | 19.85 | < **0.001** | 12.31 | < **0.001** |
| All | Location | 78.36 | < **0.001** | 39.78 | < **0.001** |
| | Stage | 68.95 | < **0.001** | 28.06 | < **0.001** |
| | Type | 0.701 | 0.403 | 76.07 | < **0.001** |
| | Location × Stage | 17.99 | < **0.001** | 11.54 | < **0.001** |
| | Location × Type | 125.6 | < **0.001** | 53.66 | < **0.001** |
| | Stage × Type | 18.42 | < **0.001** | 7.342 | < **0.001** |
| | Location × Stage × Type | 28.38 | < **0.001** | 10.78 | < **0.001** |

The significance of the influence of geographic location (indicated by different cities, Location), larval developmental stage (Stage), sample type (Type), and their interactions on alpha-diversity indices was tested. Bold $P$ values indicate significant influence ($P < 0.05$).

**Table S4.** Two-way Analysis of Variance (ANOVA) on the relative abundance of dominant prokaryotic families in larvae.

| Category | Family | Abundance rank (Descending) | Location | | Stage | | Location × Stage | |
|---|---|---|---|---|---|---|---|---|
| | | | F | *P* | F | *P* | F | *P* |
| More location-sensitive | *Vibrionaceae* | 1 | 11.88 | **< 0.001** | 6.064 | **< 0.001** | 4.425 | **< 0.001** |
| | *Flavobacteriaceae* | 2 | 28.00 | **< 0.001** | 14.20 | **< 0.001** | 6.059 | **< 0.001** |
| | *Moraxellaceae* | 3 | 37.71 | **< 0.001** | 12.52 | **< 0.001** | 12.09 | **< 0.001** |
| | *Weeksellaceae* | 4 | 17.01 | **< 0.001** | 7.804 | **< 0.001** | 3.053 | **< 0.001** |
| | *Cyclobacteriaceae* | 5 | 30.09 | **< 0.001** | 28.97 | **< 0.001** | 12.35 | **< 0.001** |
| | *Bacillales* Clade XII | 6 | 66.12 | **< 0.001** | 18.33 | **< 0.001** | 14.59 | **< 0.001** |
| | *Caryophanaceae* | 7 | 24.39 | **< 0.001** | 7.968 | **< 0.001** | 6.549 | **< 0.001** |
| | *Enterobacteriaceae* | 8 | 49.59 | **< 0.001** | 1.036 | 0.377 | 1.892 | **0.029** |
| | *Pseudomonadaceae* | 9 | 27.85 | **< 0.001** | 2.361 | 0.071 | 4.954 | **< 0.001** |
| | *Lewinellaceae* | 10 | 9.426 | **< 0.001** | 0.907 | 0.438 | 4.267 | **< 0.001** |
| | *Halieaceae* | 11 | 22.02 | **< 0.001** | 4.988 | **0.002** | 2.602 | **0.002** |
| | *Bacillaceae* | 12 | 22.26 | **< 0.001** | 0.041 | 0.989 | 1.399 | 0.156 |
| | *Streptococcaceae* | 13 | 35.49 | **< 0.001** | 3.334 | **0.019** | 8.028 | **< 0.001** |
| | *Lactobacillaceae* | 14 | 14.66 | **< 0.001** | 13.41 | **< 0.001** | 11.97 | **< 0.001** |
| | *Comamonadaceae* | 15 | 12.35 | **< 0.001** | 1.702 | 0.166 | 6.945 | **< 0.001** |
| | *Nocardioidaceae* | 16 | 46.63 | **< 0.001** | 4.629 | **0.003** | 12.91 | **< 0.001** |
| | *Micrococcaceae* | 17 | 287.4 | **< 0.001** | 6.287 | **< 0.001** | 21.18 | **< 0.001** |
| | *Sphingobacteriaceae* | 18 | 612.9 | **< 0.001** | 19.20 | **< 0.001** | 56.52 | **< 0.001** |
| More stage-sensitive | *Roseobacteraceae* | 1 | 37.66 | **< 0.001** | 45.25 | **< 0.001** | 18.30 | **< 0.001** |
| | *Pseudoalteromonadaceae* | 2 | 37.95 | **< 0.001** | 184.9 | **< 0.001** | 37.18 | **< 0.001** |
| | *Alteromonadaceae* | 3 | 14.34 | **< 0.001** | 20.75 | **< 0.001** | 10.75 | **< 0.001** |
| | *Paracoccaceae* | 4 | 5.725 | **< 0.001** | 10.64 | **< 0.001** | 4.433 | **< 0.001** |
| | *Oceanospirillaceae* | 5 | 30.44 | **< 0.001** | 65.73 | **< 0.001** | 7.733 | **< 0.001** |
| | *Erythrobacteraceae* | 6 | 4.070 | **< 0.001** | 16.56 | **< 0.001** | 3.585 | **< 0.001** |
| | *Colwelliaceae* | 7 | 21.27 | **< 0.001** | 49.82 | **< 0.001** | 11.08 | **< 0.001** |
| | *Crocinitomicaceae* | 8 | 14.44 | **< 0.001** | 31.56 | **< 0.001** | 8.000 | **< 0.001** |
| | *Spirosomaceae* | 9 | 2.987 | **0.007** | 4.382 | **0.005** | 2.340 | **0.005** |
| | *Halomonadaceae* | 10 | 2.807 | **0.011** | 12.73 | **< 0.001** | 8.049 | **< 0.001** |
| | *Piscirickettsiaceae* | 11 | 1.048 | 0.393 | 11.07 | **< 0.001** | 3.327 | **< 0.001** |
| | *Sphingomonadaceae* | 12 | 6.251 | **< 0.001** | 17.68 | **< 0.001** | 9.779 | **< 0.001** |

The significance of the influence of geographic location (indicated by different cities, Location), larval developmental stage (Stage), and their interaction on the relative abundance of the dominant families in larval prokaryotic communities (average relative abundance ≥ 2% at least at one developmental stage in one city) was tested. Bold *P* values indicate significant influence ($P < 0.05$).

**Supplementary Reference**

Allaire, J.J., Ellis, P., Gandrud, C., Kuo, K., Lewis, B.W., Owen, J., Russell, K., Rogers, J., Sese, C., Yetman, C.J., 2025. networkD3: D3 JavaScript Network Graphs from R, version 0.4.1 [software]. https://christophergandrud.github.io/networkD3/.

Apprill, A., McNally, S., Parsons, R., Weber, L., 2015. Minor revision to V4 region SSU rRNA 806R gene primer greatly increases detection of SAR11 bacterioplankton. Aquat. Microb. Ecol. 75, 129-137. https://doi.org/10.3354/ame01753.

Becker, R.A., Wilks, A.R., Brownrigg, R., Minka, T.P., Deckmyn, A., 2025. maps: Draw Geographical Maps, version 3.4.3 [software]. https://CRAN.R-project.org/package=maps.

Duan, Y., Tang, Y., Huang, J., Zhang, J., Lin, H., Jiang, S., Wang, R., Wang, G., 2020. Changes in the microbial community of *Litopenaeus vannamei* larvae and rearing water during different growth stages after disinfection treatment of hatchery water. J. Microbiol. 58, 741-749. https://doi.org/10.1007/s12275-020-0053-0.

Fox, J., Weisberg, S., 2019. An R Companion to Applied Regression, version 3.1-2 [software]. https://www.john-fox.ca/Companion/.

Hijmans, R.J., Williams, E., Vennes, C., Hijmans, M.R.J., 2026. geosphere: Spherical Trigonometry, version 1.6-8 [software]. https://CRAN.R-project.org/package=geosphere.

Kassambara, A., 2025. statix: Pipe-Friendly Framework for Basic Statistical Tests, version 0.7.3 [software]. https://CRAN.R-project.org/package=rstatix.

McMurdie, P.J., Holmes, S., 2013. phyloseq: an R package for reproducible interactive analysis and graphics of microbiome census data. PLoS One 8 (4), e61217. https://doi.org/10.1371/journal.pone.0061217.

Parada, A.E., Needham, D.M., Fuhrman, J.A., 2016. Every base matters: assessing small subunit rRNA primers for marine microbiomes with mock communities, time series and global field samples. Environ. Microbiol. 18 (5), 1403-1414. https://doi.org/10.1111/1462-2920.13023.

Reyes, G., Andrade, B., Betancourt, I., Panchana, F., Solorzano, R., Preciado, C., Sorroza, L., Trujillo, L.E., Bayot, B., 2023. Microbial signature profiles of *Penaeus vannamei* larvae in low-survival hatchery tanks affected by vibriosis. PeerJ 11, e15795. https://doi.org/10.7717/peerj.15795.

Reyes, G., Betancourt, I., Andrade, B., Panchana, F., Roman, R., Sorroza, L., Trujillo, L.E., Bayot, B., 2022. Microbiome of *Penaeus vannamei* larvae and potential biomarkers associated with high and low survival in shrimp hatchery tanks affected by acute hepatopancreatic necrosis disease. Front. Microbiol. 13, 838640. https://doi.org/10.3389/fmicb.2022.838640.

Thangaraj, S.K., Nathamuni, S.P., Katneni, V.K., Jangam, A.K., Avunje, S., Thulasi, D.N., Grover, M., Angel, J.R.J., Shekhar, M.S., 2023. Microbial communities associated with Zoea-2 syndrome and White feces syndrome in *P. vannamei* farming. Front. Mar. Sci. 10, 1120004. https://doi.org/10.3389/fmars.2023.1120004.

Wang, H., Huang, J., Wang, P., Li, T., 2020. Insights into the microbiota of larval and postlarval Pacific white shrimp (*Penaeus vannamei*) along early developmental stages: a case in pond level. Mol. Genet. Genomics 295, 1517-1528. https://doi.org/10.1007/s00438-020-01717-2.

Wang, R., Guo, Z., Tang, Y., Kuang, J., Duan, Y., Lin, H., Jiang, S., Shu, H., Huang, J., 2020. Effects on development and microbial community of shrimp *Litopenaeus vannamei* larvae with probiotics treatment. AMB Express 10, 109. https://doi.org/10.1186/s13568-020-01041-3.

Wang, Y., Wang, K., Huang, L., Dong, P., Wang, S., Chen, H., Lu, Z., Hou, D., Zhang, D., 2020. Fine-scale succession patterns and assembly mechanisms of bacterial community of *Litopenaeus vannamei* larvae across the developmental cycle. Microbiome 8, 106. https://doi.org/10.1186/s40168-020-00879-w.

Xue, M., He, Y., Chen, D., Wang, L., Liang, H., Liu, J., Wen, C.-Q., 2020. Temporal dynamics of aquatic microbiota and their correlation with environmental factors during larviculture of the shrimp *Litopenaeus vannamei*. Aquaculture 529, 735605. https://doi.org/10.1016/j.aquaculture.2020.735605.

Xue, M., Wu, L., He, Y., Liang, H., Wen, C., 2018. Biases during DNA extraction affect characterization of the microbiota associated with larvae of the Pacific white shrimp, *Litopenaeus vannamei*. PeerJ 6, e5257. https://doi.org/10.7717/peerj.5257.

Yan, M., Zhang, X., Hu, L., Huang, X., Zhou, Q., Zeng, G., Zhang, J., Xiao, G., Chai, X., Chen, J., 2020. Bacterial community dynamics during nursery rearing of Pacific white shrimp (*Litopenaeus vannamei*) revealed via high-throughput sequencing. Indian J. Microbiol. 60, 214-221. https://doi.org/10.1007/s12088-019-00853-7.

Zhang, F., Chen, H., Yu, J., Chen, R., Zhang, D., Chen, C., Wang, K., 2025. Source tracking of larval bacterial community of Pacific white shrimp across the developmental cycle: ecological insights for microbial management and pathogen prevention in larviculture. Aquaculture 609, 742800. https://doi.org/10.1016/j.aquaculture.2025.742800.
